# A differential-trend-subtype strategy for integrated serum proteomics and N-glycoproteomics reveals ovarian cancer-associated remodelling

**DOI:** 10.64898/2026.11.17.767436

**Authors:** Xiaoyu Hu, Tianqing Yan, Renquan Lu, Zhixin Tian

## Abstract

Protein glycosylation is increasingly implicated in cancer, but its macro- and microheterogeneity complicate cohort-scale resolution of disease-associated remodeling. Here we performed matched peptide-level proteomic (PEP) and intact N-glycopeptide (NGP) profiling of serum from 100 participants, comprising 23 healthy controls, 27 individuals with benign ovarian disease and 50 patients with ovarian cancer. We identified 12,634 intact NGPs with plausible linkages spanning 6,127 protein-site glycosites and 4,438 glycoproteins. We developed a Differential-Trend-Subtype (DTS) target-prioritization framework integrating ovarian cancer versus non-malignant differences, cross-sectional healthy-benign-cancer trends and within-cancer subtype effects. Independent PEP and NGP clustering supported two aligned molecular states, assigning 47 of 50 cancer samples concordantly. Among 171 subtype-associated NGPs, cluster 1-high (C1-high) features were more frequently fucosylated, whereas cluster 2-high (C2-high) features were more frequently sialylated. DTS strategy prioritized a more informative pool of candidate features. Under leakage-controlled nested resampling, the best exploratory diagnostic panel-comprising PON1, HP and C9 peptide features together with an IGHM N209 N-glycopeptide-achieved a ROC-AUC of 0.843 ± 0.026, whereas the best subtype panel-comprising C2 N621 and IGHG1 N180 N-glycopeptides together with SERPINA3-achieved a ROC-AUC of 0.997 ± 0.005. These findings demonstrate the utility of DTS for prioritizing candidate markers and reveal coordinated protein and site-specific N-glycan remodelling with potential relevance to ovarian cancer diagnosis and molecular stratification.

## 1. Introduction

Glycosylation is a non-template-driven post-translational modification (PTM) that attaches structurally diverse glycans to specific residues, expanding protein diversity. N-glycans regulate protein folding, trafficking, receptor organization, cell recognition and immune regulation, with biosynthesis dictated by the abundance and localization of glycosyltransferases, glycosidases and nucleotide-glycan donors(Moremen, Tiemeyer, & Nairn, 2012; Schjoldager, Narimatsu, Joshi, & Clausen, 2020). Because an individual glycosite can carry multiple glycan structures, these changes manifest as both macroheterogeneity (glycosylation across multiple glycosites per protein) and microheterogeneity (multiple glycosylation per glycosite at certain stoichiometry). Glycomics profiling of glycans is powerful for defining overall carbohydrate composition, but it loses the linkage between a glycan and its carrier protein and glycosite. In contrast, glycoproteomics profiling of intact glycopeptides preserves the relationship among protein, peptide, glycosite and glycan and is therefore increasingly preferred for glycoprotein analysis(Bagdonaite et al., 2022; Kawahara et al., 2021).

Recent advances in enrichment method, tandem MS fragmentation and glycoinformatics have made large-scale intact N-glycoproteomics increasingly feasible. Site-specific glycoinformatics tools of pGlyco3(Zeng, Cao, Liu, He, & Yang, 2021), Glyco-decipher(Z. Fang et al., 2022), Fragpipe(Polasky, Yu, Teo, & Nesvizhskii, 2020), StrucGP(Shen et al., 2021) accelerate joint assignment of peptide sequences, glycosites and glycan compositions; GPSeeker (commercialized as gPeptide) further resolves glycan sequence structures(Qin, Qin, & Tian, 2022; Qin, Zhang, & Tian, 2024). High-throughput serum studies show that site-specific N-glycopeptides can be quantified across clinical cohorts(Sun et al., 2023). Notably, recent studies showed that structural N-glycan changes can be extracted after accounting for protein-expression effects, separating glycan remodelling from carrier-protein abundance changes(Zhang et al., 2025). Nevertheless, serum remains analytically challenging because of its extreme concentration range, glycan structural and linkage ambiguity, co-isolation interference(P. Fang et al., 2020; Geyer et al., 2016; Malmström et al., 2016). Therefore, reliable cohort-scale serum glycoproteomics requires enrichment/fractionation and stringent quality control to mitigate the extreme dynamic range and co-isolation interference, spectrum-level evidence to distinguish linkages of N-glycopeptides, and statistical models that separate biological effects from tandem mass tag (TMT) channel effects.

Despite these analytical challenges, blood still contains proteins released from tissues together with abundant liver- and immune-related glycoproteins that report systemic responses(Pernemalm et al., 2019; Wei et al., 2021). In ovarian cancer, serum glycomics has identified alterations in high-mannose, galactosylated and differently sialylated N-glycans across healthy, benign and malignant groups, with selected glycan signatures evaluated in independent sample sets(Liu et al., 2024). Site-specific studies provide a complementary view. A serum glycoproteomic investigation of epithelial ovarian cancer quantified 653 glycopeptides and found that fucosylated tri- and tetra-antennary glycopeptides were enriched in late-stage disease, with related fucosylation patterns observed in tumour tissues(Dhar et al., 2024). A comprehensive serum glycopeptide spectral analysis detected 1,712 glycopeptides in 564 epithelial ovarian cancer cases and 1,149 controls from 13 institutions, illustrating the clinical scalability of serum-derived glycopeptide patterns(Mikami et al., 2024). Earlier work also showed increased antennarity and Lewis X-containing glycans on acute-phase proteins including haptoglobin, α1-antitrypsin and α1-antichymotrypsin in epithelial ovarian cancer(Weiz et al., 2016). Together, these observations indicate that ovarian cancer-associated serum glycosylation reflects a mixture of tumour-related carbohydrate remodelling and coordinated hepatic, inflammatory, complement and coagulation responses.

Here, we performed matched proteomic peptide (PEP) and glycoproteomic intact N-glycopeptide (NGP) profiling of a 100-sample serum cohort comprising healthy controls (HC), subjects with benign disease (Ben) and patients with ovarian cancer (OV). Using quantitative TMT-labeling technology and a high-throughput tandem mass spectrometry detection workflow, we constructed a glycoprotein landscape that covered a serum concentration range spanning up to 8 orders of magnitude. We introduce DTS, which is a three-axis analysis framework that integrates differential (D), healthy-benign-ovarian cancer trend (T), and within-ovarian-cancer subtype (S) analyses, enabling us to resolve whether disease-associated abundance changes are coupled with direction-specific preferences in glycan type, core type, fucosylation, sialylation, and other glycosylation features. Matched proteomic data were further used to distinguish NGP signals that tracked carrier-protein abundance from a smaller residual glycosylation component. Finally, we evaluated whether the combined evidence could prioritize compact candidate panels under fully nested resampling. This design positions N-glycosylation as a site-specific disease-associated phenotype and provides a framework for connecting serum carbohydrate remodelling with ovarian-cancer heterogeneity.

## 2. Materials and methods

### 2.1 Study cohort and experimental design

Serum samples were obtained from 23 healthy controls (HC), 27 participants with benign ovarian disease (Ben) and 50 participants with ovarian cancer (OV). Samples were collected at Fudan University Shanghai Cancer Center under a protocol approved by the ethics committee of Fudan University Shanghai Cancer Center with written informed consent. Clinical characteristics are summarized in **Supplementary Data 1**.

Equal 2-µL aliquots from all 100 participants were pooled as a common reference. Samples were randomized across TMT 11-plex sets, each containing ten individuals and the reference in the reporter channel 126.

### 2.2 Sample preparation, TMT labelling and intact N-glycopeptide enrichment

Proteins from 20 µL of serum were denatured in 8 M urea for 1 h and diluted with 50 mM ammonium bicarbonate (ABC). Concentration was measured by bicinchoninic acid (BCA) assay. One milligram of protein was reduced with 10 mM dithiothreitol (DTT) at 55 °C for 20 min, alkylated with 20 mM iodoacetamide for 30 min in the dark and quenched with 10 mM DTT. Samples were digested overnight at 37 °C with trypsin at the enzyme-protein-ratio of 1:50 (w/w), desalted on C18 solid phase extraction columns (Phenomenex Jupiter, 15 µm, 300 Å) and vacuum dried.

Peptides were dissolved in 100 mM triethylammonium bicarbonate (TEAB) and labelled for 1 h with TMT 11-plex reagent in anhydrous acetonitrile (ACN) at a 1:1 (w/w) peptide-reagent-ratio. Reactions were quenched with 5% hydroxylamine for 15 min before pooling and desalting. Intact N-glycopeptides were enriched on in-house ZIC-HILIC columns (Merck; 5 µm, 200 Å) conditioned three times with 300 µL of 80% ACN containing 1% trifluoroacetic acid (TFA). Peptides were reconstituted in this solution, loaded five times and washed. Flow-through and wash fractions were combined for peptide analysis; retained N-glycopeptides were eluted with 300 µL of 0.1% TFA followed by 200 µL of 50 mM ABC, pooled and dried.

### 2.3 High-pH fractionation and LC-MS/MS acquisition

Approximately 100 µg of enriched N-glycopeptides per TMT set were fractionated with a Pierce High pH Reversed-Phase Peptide Fractionation Kit (Thermo Fisher Scientific; No. 84868). Samples in 300 µL of 0.1% TFA were loaded and centrifuged at 3,000 × g for 2 min. After a wash with 5% ACN and 0.1% triethylamine, eight fractions were eluted with 10%, 12.5%, 15%, 17.5%, 20%, 22.5%, 25% or 50% ACN containing 0.1% triethylamine, dried separately and analysed by liquid chromatography-tandem mass spectrometry (LC-MS/MS).

TMT-labelled peptides were loaded on an Ultimate 3000 nano-LC coupled to a Q Exactive mass spectrometer (Thermo Fisher Scientific). A 65-cm analytical column (360 µm outer × 75 µm inner diameter) and 5-cm trap column (360 µm outer × 200 µm inner diameter) were packed with Phenomenex Jupiter C18 resin (5 µm, 300 Å). Mobile phases A and B were 0.2% formic acid in water and in 80% ACN, respectively, at 300 nL/min. Peptides were loaded at 2-12% B for 10 min, separated at 12-45% B over 190 min, ramped to 95% B over 10 min, held for 5 min, returned to 2% over 10 min and re-equilibrated for 10 min.

Peptides were ionised at 2.8 kV with the ion transfer capillary at 250 °C. Full spectra (*m/z* 350-1,800) were acquired at 70,000 resolution, an automatic gain control target of 2 × 10^5^ and a 50-ms maximum injection time. The TOP 20 precursors underwent higher energy collisional dissociation at a normalized collision energy of 30%. MS2 spectra were acquired at 17,500 resolution with a 1.4 *m/z* isolation window, a target of 5 × 10^5^ and a 250-ms maximum injection time. Singly charged ions and charge states ≥7 were excluded; dynamic exclusion was 20 s. Each TMT set was injected three times.

Intact N-glycopeptides were loaded on the same platform and columns. Samples were loaded at 2-10% B for 10 min and separated at 10-40% B over 190 min, followed by the same ramp, wash and equilibration steps. Full spectra (*m/z* 700-2,000) were acquired at 70,000 resolution, a target of 2 × 10^5^ and a 50 ms maximum injection time. The 20 most intense precursors underwent stepped higher energy collisional dissociation at 20%, 30% and 31%. MS2 spectra were acquired at 17,500 resolution with a 1.4 *m/z* isolation window, a target of 5×10^5^ and a 250 ms maximum injection time.

### 2.4 Peptide and intact N-glycopeptide identification

Peptide raw files were processed with FragPipe v22.0 and MSFragger v4.1 against the UniProt Homo sapiens reference proteome downloaded on 16 April 2024 (20,431 sequences). Searches used strict trypsin specificity, up to two missed cleavages and 20 ppm precursor and fragment tolerances. “TMT-10” workflow was used. Carbamidomethylated cysteine and TMT labelled lysine were fixed; oxidation on methionine, protein N-terminal acetylation and TMT labelled peptide N termini were variable. Peptide spectrum matches (PSMs) and proteins were controlled at 1% false discovery (FDR) by a target-decoy strategy. Reporter ions were quantified with IonQuant v1.10.27 and TMT-Integrator v5.0.9.

Intact N-glycopeptides were identified with gPeptide against the same proteome and a customised database containing 5,820 N-glycan linkages. Searches allowed up to two missed tryptic cleavages and used precursor and fragment ion tolerances of 10 and 20 ppm, respectively. Identifications required experimental observation of all isotopic peaks with relative abundance ≥40%, isotope peak abundance deviation ≤50%, at least five matched peptide backbone product ions and a Y1 ion ranked among the top four candidates. Spectrum FDR was controlled at 1% using a target-decoy strategy. Assignments within each dataset and spectrum index were ranked by the new pg score; exact top score ties between feature identities were excluded. An NGP feature was defined by protein accession, peptide sequence, N-glycosite, N-glycan composition and N-glycan linkage.

### 2.5 Data processing and quality control

For each retained PSM of PEP, the log_2_ ratio of every biological reporter channel to the pooled 126 channel was calculated. Ratios were median aggregated by feature and channel within each technical replicate and then across the three replicates with equal weight. For NGP, reporter to 126 ratios from uniquely assigned spectra were log_2_ transformed and median aggregated across retained GPSMs and fractions for each feature and TMT set. The resulting 100 sample PEP and NGP abundance matrices were centred by subtracting each sample median.

Features detected in more than 70% of samples formed the 70% coverage matrix used for multivariate and subtype analysis. Missing values in full cohort descriptive analysis were imputed by k nearest neighbours (k=10). In resampled model evaluation, coverage filtering, imputation, nuisance correction and scaling were estimated from training data only. Pairwise complete Pearson correlations were calculated from the 512 non-imputed NGP features with more than 70% coverage. HC and Ben samples were ordered together by average linkage clustering and OV samples independently. Within-group concordance was the mean correlation with other samples in the same group.

Unique N-glycopeptide sequences were aligned from -10 to +10 around the glycosite to calculate position specific amino acid frequencies. Glycans were annotated by type, core, antenna count, sialylation and fucosylation; the latter was classified as absent, core fucose (CF), branch fucose (BF) or both. Proteins and N-glycoproteins were matched to Human Protein Atlas (https://www.proteinatlas.org/) blood concentrations by UniProt accession or unambiguous gene symbol, and analysis was restricted to matched proteins. Positive PSM and GPSM reporter signals were summed by protein within replicate and averaged across replicates and clinical groups.

### 2.6 Differential and trend analysis

Differential (D) and trend (T) analysis was performed separately for PEP and NGP with limma v3.64.3. D features were identified by comparing OV with HC and Ben combined in models containing diagnostic group. T was modelled by coding the cross-sectional HC-Ben-OV disease states as 0, 1 and 2, with TMT set and reporter channel as covariates; adjusted group means were also required to be monotonic. Robust empirical Bayes moderation with intensity trend estimation was applied. Features were retained at a Benjamini-Hochberg FDR below 0.05 and an absolute fold change of at least 1.3. The D effect was the log_2_ fold change for OV versus HC and Ben combined, and the T effect was the adjusted OV minus HC difference.

Features were classified as D&T, D only, T only or neither. D and T endpoint effects were compared by Pearson correlation, and concordance was the percentage of D&T features with matching signs. Shared D&T NGPs were partitioned by the maximum silhouette coefficient across k=2-6. Glycan traits were compared across expression pattern groups by chi-squared tests with 5,000 permutations and Benjamini-Hochberg correction. Each displayed feature had at least one spectrum containing at least five matched peptide-backbone product ions, single-site diagnostic evidence and structure-diagnostic ions in the same PSM. Within these evidence constraints, features were selected to retain effect direction and glycan-structure diversity; this display prioritization did not define an additional identification FDR or localization probability.

Features were collapsed to unique protein accessions before separate STRING v12.5 enrichment analysis of the PEP-D, PEP-T, NGP-D and NGP-T sets against the corresponding detectable protein universe. Terms with a STRING FDR below 0.05 were retained.

### 2.7 Subtype and heterogeneity analysis

OV samples were clustered independently in the PEP and NGP layers with ConsensusClusterPlus v1.72.0. The 70% coverage matrices were imputed by k nearest neighbours, corrected for TMT set and ranked by median absolute deviation. The top 200 features were used for the primary analysis with Pearson distance, average linkage, 500 resamples and item and feature sampling proportions of 0.8. Solutions from k=2 to 8 and sets of 100, 200 and 300 features were compared by ambiguous clustering, within-cluster consensus, silhouette width, minimum cluster size and cross-omic adjusted Rand index. For k=2, C1 and C2 were oriented as the subtypes with relatively lower and higher disease signals, respectively. Cross-omic agreement was summarized by agreement, adjusted Rand index and normalized mutual information.

Subtype associated (S) effects were estimated among the 50 OV samples with limma models containing subtype, TMT set and reporter channel; the effect was C2 minus C1. S features met a Benjamini-Hochberg FDR below 0.05 and an absolute fold change of at least 1.3. Clinical and technical covariates were assessed using exact tests appropriate to the structure of the contingency tables. Potential confounding by disease stage was evaluated through stage-adjusted and stage-restricted sensitivity analyses. Directional disease and subtype scores were calculated as the mean feature z-score multiplied by the sign of the corresponding D or S effect, and their association among OV samples was assessed by Spearman correlation.

For NGPs with matched peptide carrier abundance, PEP protein abundance was the median of the corresponding peptide features. Raw NGP abundance, NGP minus PEP abundance and residuals from NGP-on-PEP regression were compared in a common 275-feature universe. Features detected in at least 70% of samples in all three layers were retained and imputed by k nearest neighbours (k=10). Each layer was tested in a limma C2 versus C1 model containing TMT set and reporter channel, anchored to the PEP-derived state assignment. The retention criteria were a Benjamini-Hochberg FDR below 0.05 and an absolute fold change of at least 1.5.

### 2.8 Model training and evaluation

Diagnostic models distinguished OV from HC and Ben combined using five repeats of five stratified outer folds shared across comparisons. Coverage filtering, median imputation, TMT set correction, scaling, feature ranking, selection and hyperparameter tuning were fitted within each outer training fold; reporter channel was not removed separately. For DTS axis, percentile ranked absolute effect size and -log_10_(FDR) were combined by geometric mean, with non-monotonic T evidence set to zero; the DTS score was the geometric mean across axes. Panels of 5, 10 and 20 features were evaluated by ridge logistic regression, and stability by the mean pairwise Jaccard index.

At the ten-feature budget, D only and continuous DTS rankings were compared using ridge and elastic net logistic regression, linear and radial basis function support vector machines, random forest and gradient boosting. Hyperparameters were selected by fourfold inner cross-validation. Outer test performance was summarized by the area under the receiver operating characteristic curve (ROC-AUC); precision-recall AUC, balanced accuracy and Brier score were secondary measures.

Fully nested OV state rediscovery used ten repeats of five folds, approximately stratified by TMT set and generated without subtype labels. Each outer training fold applied 70% coverage filtering, training-only k nearest-neighbour imputation, training-estimated set correction, top-200 median absolute deviation selection, scaling and k=2 consensus clustering with 500 resamples. Held-out samples were assigned to the nearest training centroid by Pearson correlation distance. Orientation used the training fold disease direction; fixed full cohort labels were used only to assess concordance after prediction.

Candidate panels were evaluated using five repeats of five outer folds and three repeated fourfold inner splits. Candidate subsets and elastic net hyperparameters were selected within inner resampling, and the selected configuration was fitted to the complete outer training fold before prediction. Repeat aggregated ROC curves were calculated from outer test predictions.

### 2.9 Statistical analysis

Unless otherwise specified, tests were two sided and multiplicity was controlled by the Benjamini-Hochberg procedure. A FDR below 0.05 was considered significant. Analysis was performed in R v4.5.1 and Python; software-specific versions are reported above.

## 3. Results

### 3.1 Intact N-glycopeptide identification displayed a broad and heterogeneous serum glycosylation landscape

The experimental and analytical workflow is summarized in **Figure 1A**. Quality control for PEP and NGP data showed consistent feature recovery and high replicate agreement across both molecular layers (**Supplementary Fig. 1**).

**Figure 1.**
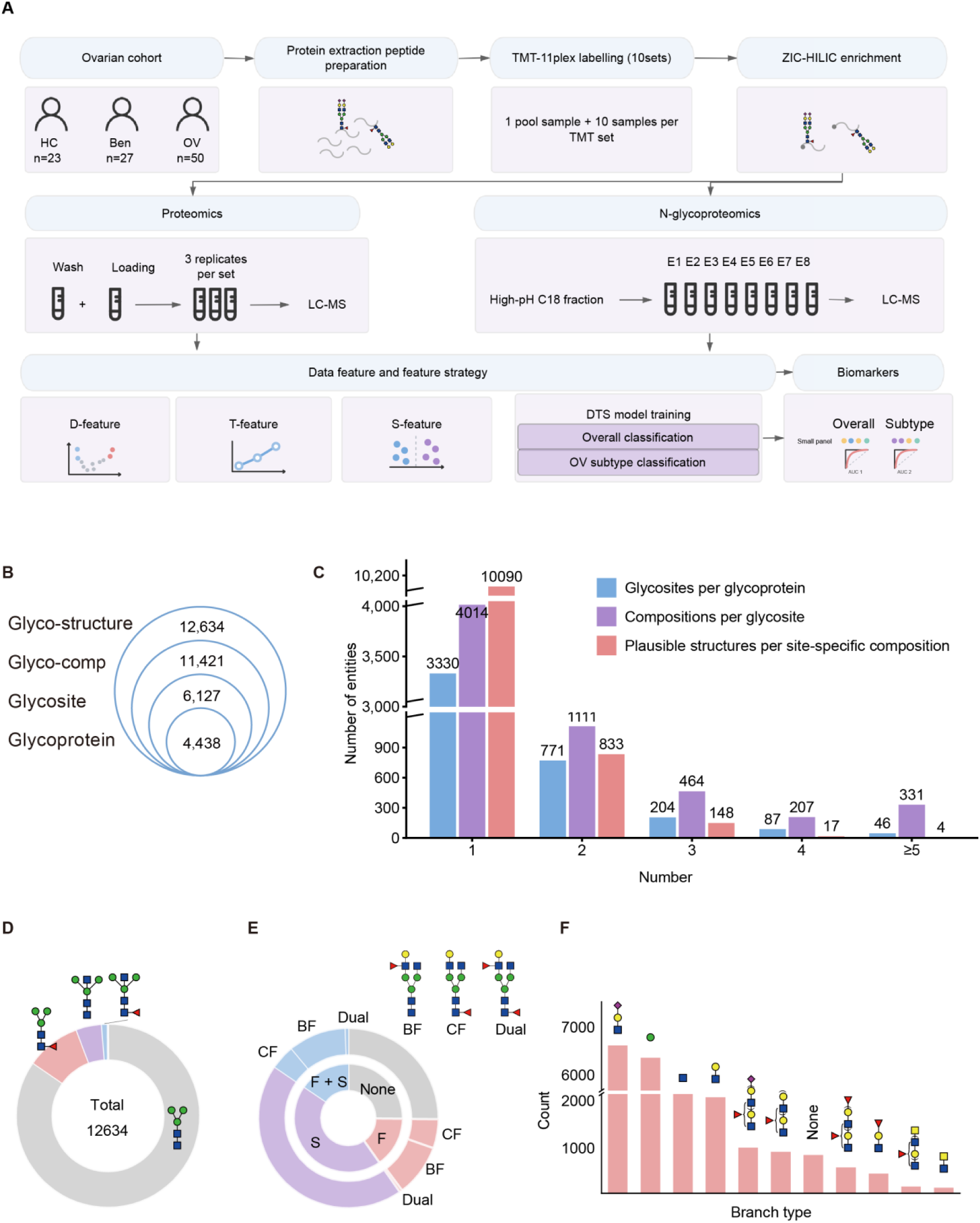
Study design and qualitative landscape of the serum peptide and intact N-glycopeptide data. (A) Experimental and analytical workflow for 23 healthy controls (HC), 27 participants with benign ovarian disease (Ben) and 50 participants with ovarian cancer (OV). Peptide (PEP) and intact N-glycopeptide (NGP) data were integrated through differential (D), trend (T) and subtype (S) analysis. (B) Qualitative identifications obtained in this study. (C) Distributions of glycosites per glycoprotein, glycan compositions per glycosite and plausible linkage structures per site-specific composition. (D) Distribution of glycan core types. (E) Cross-classification of fucosylation (F) and sialylation (S); CF, core fucosylation; BF, branch fucosylation; dual, CF and BF. (F) Counts of branch motifs across glycan antennae.

Across the 100-sample serum cohort, we identified 12,634 intact N-glycopeptides (NGPs), defined as protein-peptide-glycosite-composition-linkage features. These represented 11,421 distinct protein-peptide-glycosite-composition entities, 6,581 protein-peptide-glycosite backbones and 4,438 glycoproteins (**Fig. 1B**). NGPs and PEPs were provided in **Supplementary Data 2** and **Supplementary Data 3**, respectively. Collapsing peptide redundancy yielded 6,127 unique protein-site glycosites. The heterogeneity of glycosylation was also diverse.(**Fig. 1C**). Most glycoproteins carried one N-glycosite (3,330 of 4,438; 75.0%), 771 (17.4%) carried two, 204 (4.6%) carried three, 87 (2.0%) carried four and 46 (1.0%) carried at least five, with a maximum of 22 sites. Site-level composition was similarly concentrated: 4,014 of 6,127 glycosites (65.5%) bore one glycan composition, 1,111 (18.1%) bore two, 464 (7.6%) bore three, 207 (3.4%) bore four and 331 (5.4%) bore at least five. Linkage resolution added a further isomerism layer. Of 11,092 site-specific compositions, 10,090 (91.0%) resolved to one plausible linkage, 833 (7.5%) to two, 148 (1.3%) to three, 17 (0.2%) to four and four (0.04%) to at least five. Linkage isomerism with monosaccharide sequence resolution therefore affected 9.0% of site-specific compositions. Moreover, 9,282 of the 12,634 unique intact N-glycopeptide IDs (73.5%) were supported by at least one retained spectrum containing a structure-diagnostic glycan fragment ion, providing fragment-level support for the assigned monosaccharide sequence.

Glycans were predominantly complex (6,314; 50.0%) or hybrid (4,527; 35.8%), with a smaller high-mannose fraction (1,793; 14.2%). Across the 12,634 NGPs, most glycans carried a common core (10,691; 84.6%); the remainder were core-fucosylated (1,212; 9.6%), carried a bisecting GlcNAc (571; 4.5%) or both (160; 1.3%) (**Fig. 1D**). Across all the NGPs, 5,582 (44.2%) were sialylated only, 1,910 (15.1%) were fucosylated only, 1,958 (15.5%) carried both and 3,184 (25.2%) carried neither (**Fig. 1E**). Branch fucosylation outnumbered core fucosylation among both the fucosylated-only (1,188 versus 661) and doubly modified (1,308 versus 560) classes. Terminal and branch motifs summed across antennae (20,959 counts) were dominated by sialic-acid (6,577) and mannose (6,320) termini, followed by terminal GlcNAc (2,157) and terminal galactose (2,029); fucosylated termini contributed 2,530 (sialic-acid 963, galactose 876, fucose 548, di-HexNAc 143), with fucose (419) and di-HexNAc (118) termini (**Fig. 1F**). The identified sequences recapitulated the canonical N-glycosylation sequence N-X-S/T (X ≠ P). Aligning the glycopeptide sequences across positions −10 to +10 flanking the modified asparagine, Asn occupied position 0, Ser/Thr were enriched at position +2 and Pro was depleted at position +1 (**Supplementary Fig. 2**).

### 3.2 Deep identification for glycoproteomics

Site-specific mapping illustrated microheterogeneity within individual glycoproteins (**Fig. 2A**). Attractin (ATRN, O75882) carried high-mannose, complex and hybrid glycans across nine observed N-glycosites, and several site-specific compositions resolved to multiple plausible linkages. Regarding the aspect of micro-heterogeneity, there is a rich composition of glycan groups at the N46 of the IGHM (**Fig. 2B**). Out of the 759 GPSMs for N46, 567 contain site-diagnostic ions, and 622 contain structural-diagnostic ions, which can be used to determine the local structure.

**Figure 2.**
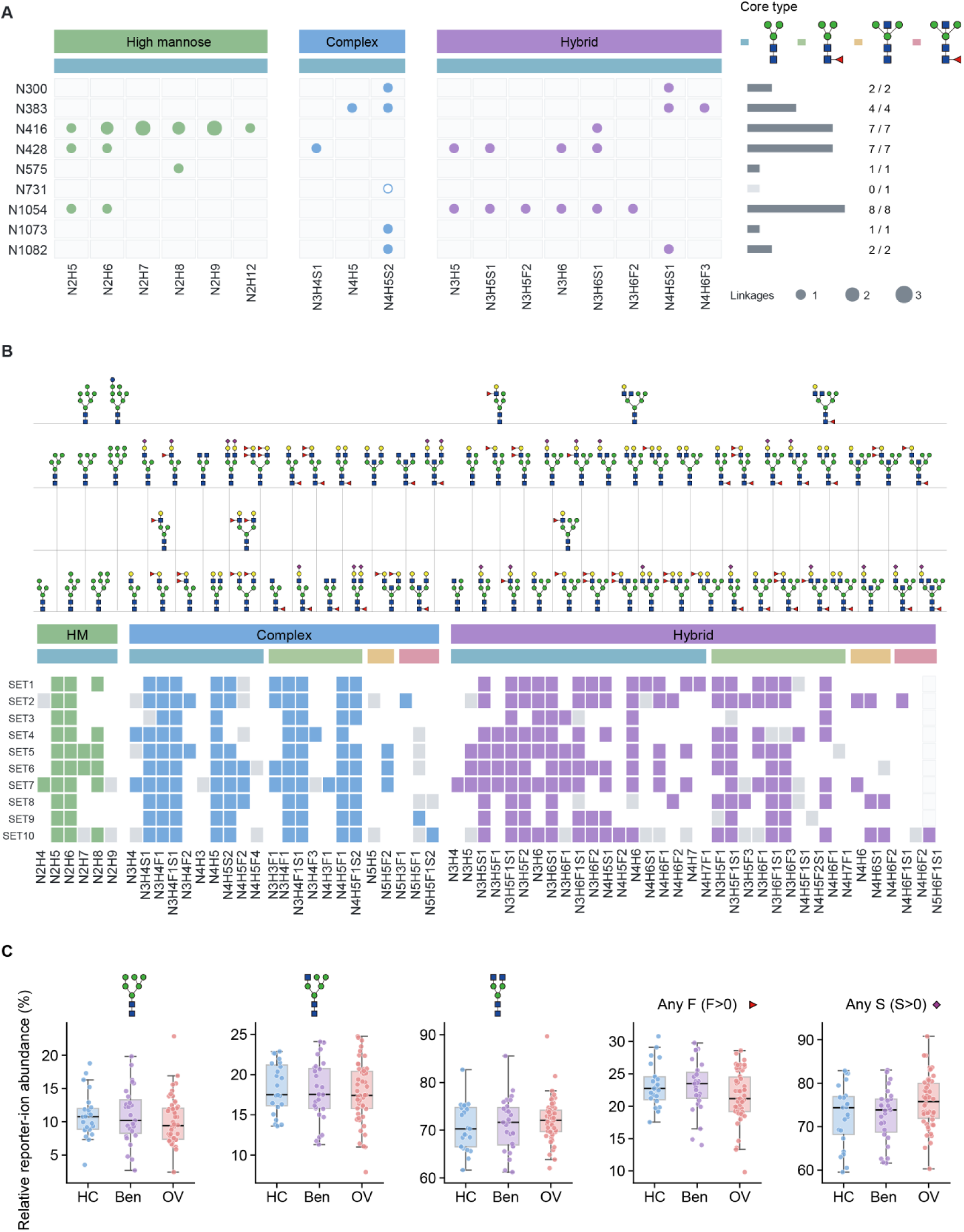
Site-specific N-glycan microheterogeneity and recurrence across TMT sets. (A) Glycan compositions detected at nine N-glycosites of attractin (ATRN; O75882), grouped as high-mannose (HM), complex or hybrid types. (B) Detection of representative glycan compositions across SET1-SET10, grouped by glycan class and core modification.

To benchmark the abundance range captured by the two workflows, we matched qualitatively identified proteins to mass-spectrometry-based blood concentrations in the Human Protein Atlas. Reference concentrations were available for 1,104 of 4,438 NGP proteins and 169 of 264 PEP proteins. NGP covered 2.9×10^3^-4.2×10^11^ pg·L^-1^, extending approximately 21-fold below the lowest PEP-matched concentration and showing a lower median concentration than PEP (6.15×10^5^ versus 7.70×10^9^ pg·L^-1^; **Supplementary Fig. 3A-C**). Reporter-signal profiles were also less top-heavy: the ten most abundant proteins accounted for 15.1-18.8% of the NGP signal across HC, Ben and OV, in contrast to 89.0-90.4% of the PEP signal (**Supplementary Fig. 3D-I)**. These results indicate that NGP profiling extends the accessible serum concentration range and distributes signal across a broader protein repertoire.

### 3.3 Differential and trend analysis converges on a shared circulating signature

PCA visualized global PEP and NGP variation across HC, Ben and OV (**Supplementary Fig. 4A-D**), but HC and Ben were not well separated (**Supplementary Data 4**). We therefore defined differential (D) features as the fold-change value of OV relative to HC+Ben. Differential (D) and trend-associated (T) features were highly concordant across both molecular layers (**Supplementary Data 5**). Among NGPs, 68 of 88 D features were also identified by the trend analysis, representing 85% of the 80 T features (**Fig. 3A**). D and T features were closely correlated for PEP (r=0.965) and NGP (r=0.980), and all 99 shared PEPs and 68 shared NGPs changed in the same direction (**Fig. 3B**). Thus, the cross-sectional HC-Ben-OV trends largely recapitulated the OV versus HC/Ben contrast.

**Figure 3.**
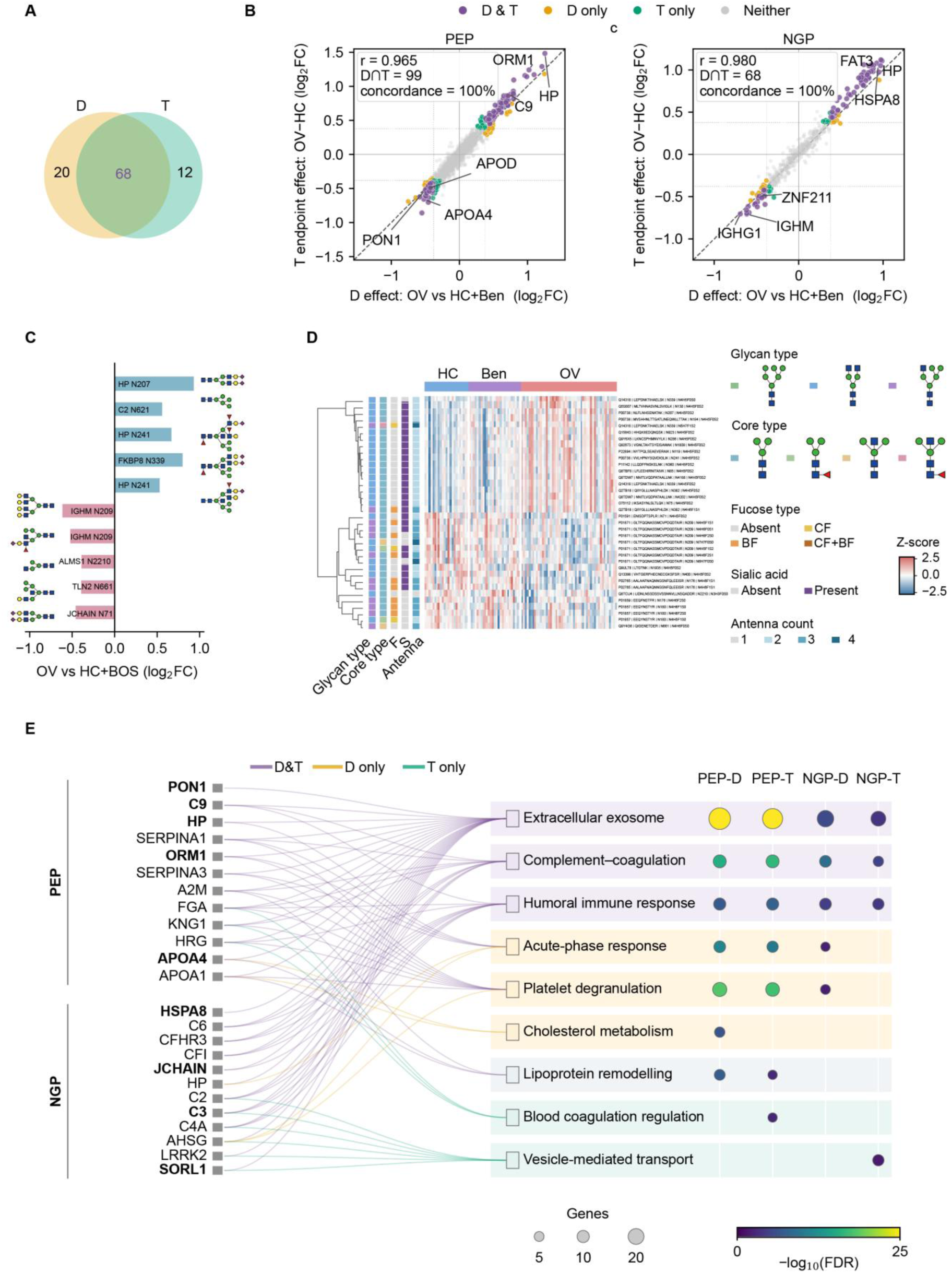
Differential and trend analysis converges on a shared circulating signature. (A) Overlap of D- and T-NGPs. (B) D effects plotted against T endpoint effects for PEP and NGP features. (C) Adjusted D effects for five OV-high and five OV-low D&T-shared glycopeptides prioritized by strict PSM evidence and glycan-structure diversity; protein, glycosite and glycan composition are indicated. (D) Row-wise z-scored abundances of representative shared NGPs across HC, Ben and OV samples, with expression and glycan annotations. (E) Integrated protein-to-pathway network and enrichment bubble matrix for the PEP-D, PEP-T, NGP-D and NGP-T sets.

Abundance profiles of representative shared NGPs resolved reciprocal OV-low and OV-high pattern groups (**Fig. 3D**). All 68 NGPS were showed in **Supplementary Figure 4E.** They were predominantly sialylated (75%), carried complex (59%) or hybrid (37%) glycans and most commonly contained an unmodified core (87%); 71% lacked fucose (**Supplementary Fig. 5A**). Among features carrying core fucose or branch fucose, 75% and 60% were OV-low, respectively, whereas 80% of sialylated features and 87% of biantennary features were OV-high (**Supplementary Fig. 5B**). Ten PSM-supported representative glycopeptides spanning both directions and structurally diverse glycan classes are shown by adjusted D effect in **Fig. 3C**. The OV-high HP N207_NLFLNHSENATAK_N4H5F0S2 recapitulated previously reported site-specific hyper-sialylation of the same haptoglobin peptide in ovarian-cancer serum(Shetty, Hafner, Shah, Nickens, & Philip, 2012). Besides, N241 of HP was also reported in previous study(Weiz et al., 2016). The IGHM N209 glycopeptide carrying the N4H5F1S1 composition matched a glycosite and composition previously reported in independent EOC cohorts(Ruhaak et al., 2016), although its disease-associated direction was opposite in the present cohort.

STRING analysis further organized these concordant signals into shared and evidence-specific functional branches (**Fig. 3E**). Extracellular exosome, complement-coagulation and humoral immune-response terms were supported across PEP-D, PEP-T, NGP-D and NGP-T. Acute-phase response and platelet degranulation were shared by PEP-D, PEP-T and NGP-D, whereas PEP additionally resolved lipid-related terms, including D-specific cholesterol metabolism and D/T-supported lipoprotein remodelling. Blood coagulation regulation was specific to PEP-T, and vesicle-mediated transport to NGP-T. Cross-study comparison at exact protein–site–composition identified 262 shared NGPs between the present serum dataset and tissue dataset (Hu et al., 2020; Pan et al., 2020), of which 38 were direction-comparable. Concordance predominated among liver/secreted and ECM/stromal units, whereas 11 of 12 complement/coagulation units were discordant; within-site direction switching at C3 N85, FN1 N528 and HPX N187 further suggested compartment-specific glycoform redistribution (**Supplementary Fig. 6)**. Together, these results define a common circulating disease-state signature with smaller layer- and criterion-specific branches. We therefore next asked whether this shared programme was uniformly expressed across OV samples or stratified into reproducible molecular states.

### 3.4 Cross-omic consensus supports two ovarian cancer molecular states

OV samples were clustered independently within each omic layer using a prespecified k=2 procedure (TOP200 median-absolute-deviation (MAD) features, Pearson distance, average linkage, 500 resamples; **Fig. 4A**). PEP and NGP returned partitions of 25/25 and 24/26 samples, respectively, and the two layers assigned the same state to 47 of 50 OV samples (**Fig. 4B**). Three discordant samples (OV051, OV083 and OV169) were marked throughout. Consensus diagnostics across cluster numbers and feature-set sizes, together with cross-omic agreement, supported the two subtypes of NGPs (**Supplementary Fig. 7A-E**).

**Figure 4.**
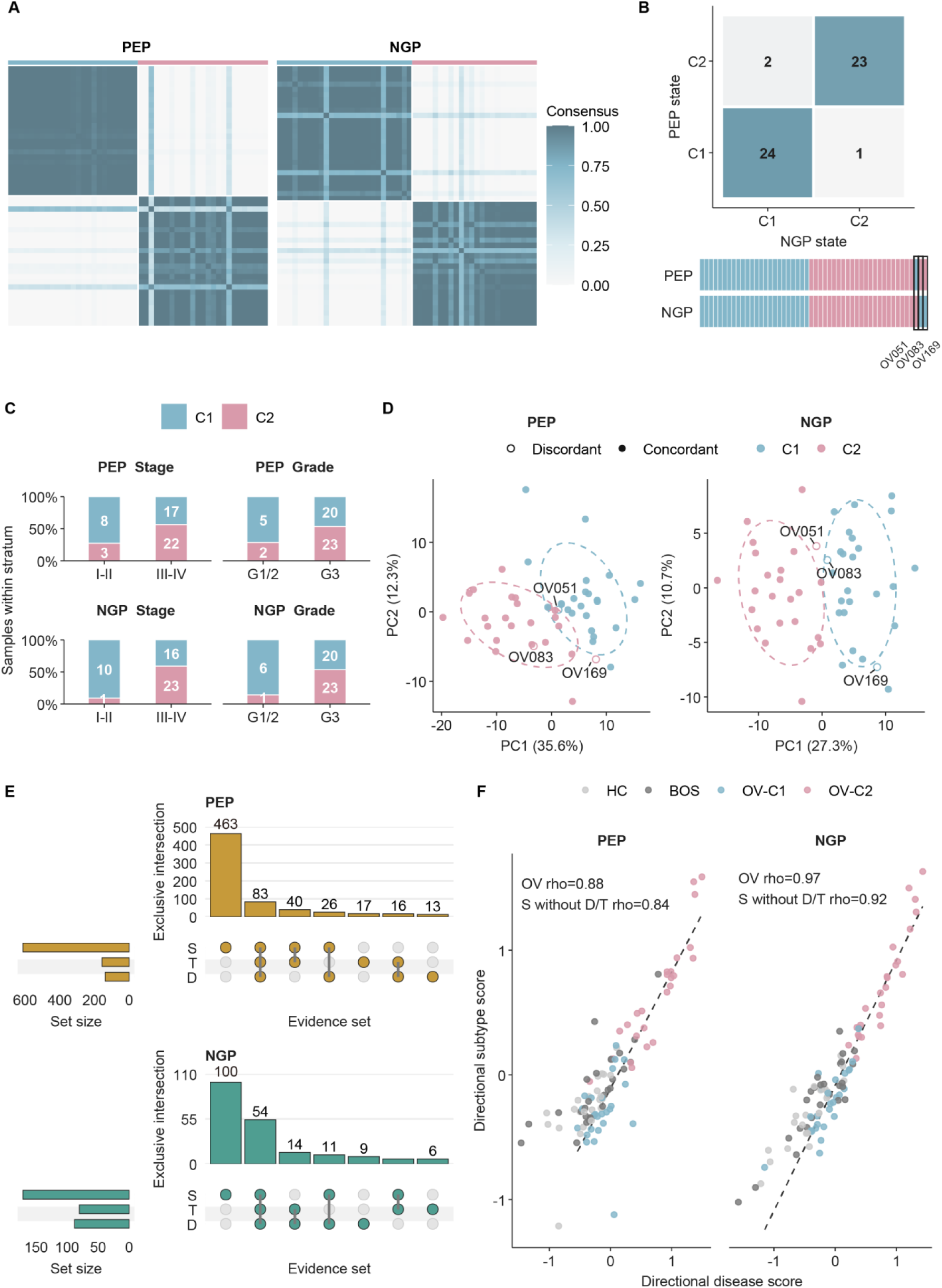
Independent peptide and glycopeptide clustering supports two ovarian cancer molecular states. (A) Consensus matrices from independently clustered PEP and NGP profiles using the prespecified k=2 procedure. (B) Cross-omic assignment matrix and sample-level tracks. (C) Distribution of stage and grade within PEP- and NGP-derived C1 and C2 states. (D) PCA of OV samples, with discordant samples labelled. (E) Exclusive intersections and total sizes of the D, T and S sets for PEP and NGP. (F) Directional disease and subtype scores across HC, Ben, OV-C1 and OV-C2 samples; ρ values are Spearman correlations among the 50 OV samples, with values after excluding D/T-linked S features shown separately.

The C1 and C2 molecular states were then audited against clinical and technical covariates (**Fig. 4C**). The NGP subtype was associated with FIGO stage (*p*=0.0051, *q*=0.040), whereas the PEP stage association was not significant (*p*=0.17). Same-cohort PCA visualized separation of the C1 and C2 states (**Fig. 4D**). Therefore, both PEP and NGP showed the expression directions of the two branches in the OV group (**Supplementary Fig. 7F**). With the original C1/C2 labels fixed, stage adjustment retained 544 of 612 PEP and 149 of 171 NGP subtype-associated features, whereas restriction to stage III-IV disease retained 317 and 108 features, respectively (**Supplementary Fig. 7G**).

At the feature level, 612 PEP and 171 NGP features passed the FC=1.3 threshold for the subtype (S) axis, and the exclusive intersections and total sizes of the differential (D), trend (T) and subtype (S) feature sets were displayed (**Fig. 4E, Supplementary Data 5**). Directional disease and subtype scores were then computed for all samples and correlated across the 50 OV samples (**Fig. 4F**). Directional disease and subtype scores were strongly correlated among OV samples in both molecular layers. The associations remained strong after excluding S features that overlapped either D or T. Thus, the C1-C2 subtype architecture was aligned with the magnitude of disease-associated molecular variation, particularly in the NGP layer, and this relationship was not explained solely by direct overlap between the D/T and S feature sets. Features shared across the differential, trend and subtype analysis retained reciprocal C1-high and C2-high cross-sectional abundance patterns (**Supplementary Fig. 7F**).

### 3.5 Coordinated proteomics and glycoproteomics define the two ovarian cancer molecular subtypes

32 representative S features comprising 16 C1-high and 16 C2-high features per heatmap from each molecular layer were displayed (**Fig. 5A**). Across the complete NGP-S set, 91 of 171 glycopeptides were C1-high and 80 were C2-high, C2-high features showed more sialylation (87.5%) and four-antenna glycans (7.5%), whereas C1-high features showed more total fucosylation (27.5%) and core fucosylation (12.1%, **Fig. 5B**). Glycan linkage distributions of C1-high and C2-high NGPs are shown in **Supplementary Fig. 8A-F.**

**Figure 5.**
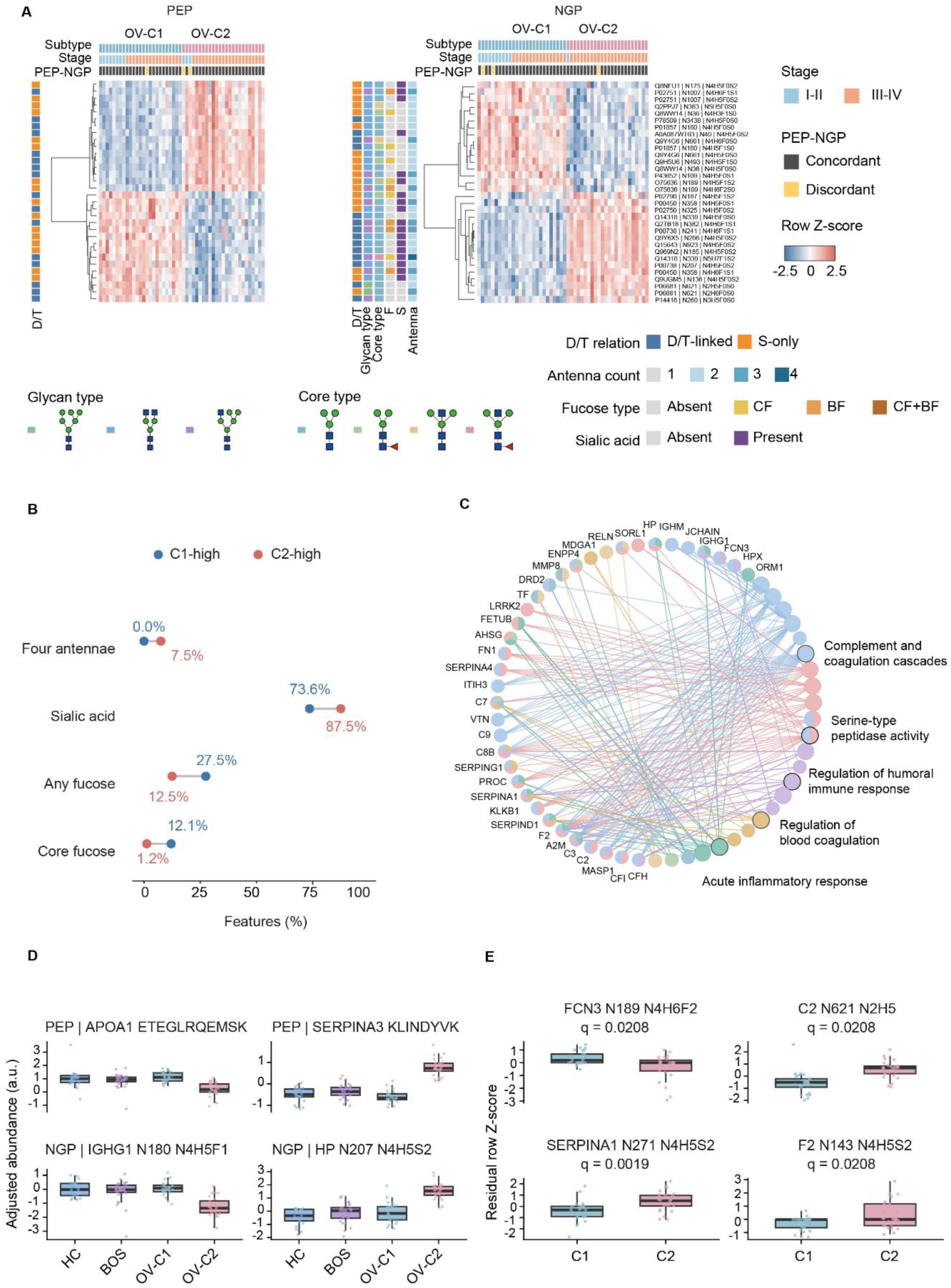
Molecular architecture of the two ovarian cancer states and residual glycosylation after carrier adjustment. (A) Row-wise z-scored PEP and NGP subtype heatmaps containing 32 representative S features per layer. (B) Percentages of tetra-antennary, sialylated, fucosylated and core-fucosylated NGP-S features among the 91 C1-high and 80 C2-high glycopeptides. (C) Cytoscape network of subtype-associated S proteins. (D) Representative PEP and NGP abundance profiles across HC, Ben, OV-C1 and OV-C2. (E) Carrier-adjusted regression residuals for four glycopeptides.

The 171 NGP-S features mapped to 108 glycoproteins. A network based on these S proteins linked the subtype-associated carriers to complement and coagulation cascades, serine-type peptidase activity, regulation of humoral immune response, regulation of blood coagulation and acute inflammatory response (**Fig. 5C**). Representative peptides and N-glycopeptids illustrated the coordinated abundance patterns across HC, Ben, OV-C1 and OV-C2 samples (**Fig. 5D**).

Because glycan-level changes can reflect carrier-protein abundance, we repeated the subtype contrast after correcting NGP abundance for matched peptide-level carrier abundance. Across the same 275 correction-eligible NGPs, using the PEP-derived C1/C2 assignment as a cross-omic anchor, 48 raw NGPs, nine NGP-minus-PEP features and four regression-residual glycopeptides met BH-FDR < 0.05 and an absolute fold change of at least 1.5. The four residual glycopeptides were FCN3_N189_N4H6F2, C2_N621_N2H5, SERPINA1_N271_N4H5S2 and F2 N143_N4H5S2 (**Fig. 5E**); three were C2-high and FCN3 was C1-high. Thus, most subtype-associated NGP signal tracked carrier-protein abundance, whereas a smaller residual glycosylation tier remained. The information of these four protein-glycosites has been reported by Uniprot, and site-diagnostic ions were also identified in this study. Fucosylation of FCN3 has also reported, even though it differs from the linkage assumed in the previous article(Jia et al., 2009). Pan et al. assigned SERPINA1_N271_N4H5S2 to IGP group 3, characterized by elevated fucosylation and sialylation, whereas Hu et al. assigned it to IG3, a fucose/sialic acid-associated group enriched in complement and coagulation pathways., representing the group with elevated levels of fucose and sialic acid. Besides, it was assigned by Hu et al. to IG3, a fucose/sialic acid-associated group enriched in complement and coagulation pathways.

### 3.6 Diagnostic and subtype biomarker discovery

Under a leakage-controlled 5 × 5 nested benchmark distinguishing OV from HC and Ben combined, single-layer peptide and intact N-glycopeptide models performed equivalently and modestly (ROC-AUC 0.726±0.024 and 0.726±0.036, respectively). We therefore asked whether fold-local integration of differential (D), cross-sectional HC-Ben-OV trend (T) and subtype (S) evidence, reconstructed from the training data only, could prioritize compact diagnostic models more effectively. At five selected features, mean ROC-AUC rose from 0.700±0.027 for D-only ranking to 0.721±0.035 with D+T and 0.728±0.021 for continuous DTS; at ten features the corresponding values were 0.711±0.023, 0.726±0.031 and 0.730±0.016, whereas at 20 features the three schemes converged (**Fig. 6A**). Feature-selection stability at ten features increased from a mean Jaccard index of 0.238 for D-only ranking to 0.389 for continuous DTS. At the ten-feature budget, continuous DTS improved mean ROC-AUC for five of six learners, including ridge logistic regression (0.711 to 0.730) and gradient boosting (0.667 to 0.722); the RBF support-vector machine was the exception (0.703 to 0.695) (**Fig. 6B**). A broader comparison of D-, T- and S-based ranking strategies across panel sizes, together with classifier-specific differences between continuous DTS and D-only ranking, is shown in **Supplementary Fig. 9A-C**.

**Figure 6.**
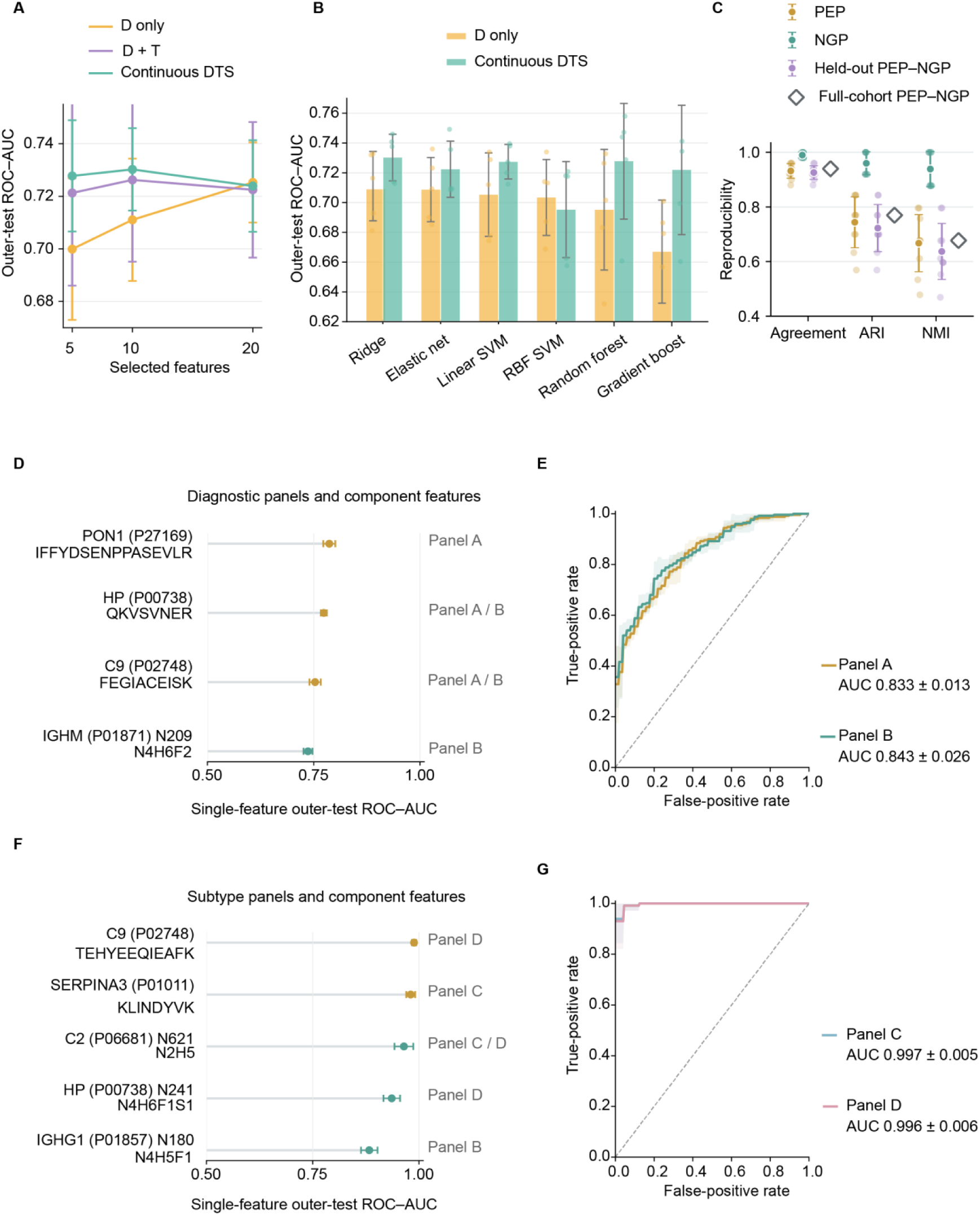
Fold-local DTS prioritization and exploratory compact biomarker panels. (A) Outer-test ROC-AUCs for D-only, D + T and continuous DTS ranking using 5, 10 or 20 selected features in leakage-controlled 5 × 5 nested cross-validation. (B) Ten-feature outer-test ROC-AUCs for D-only and continuous DTS ranking across six classifiers. (C) Reproduction and concordance of de novo k=2 subtype assignments across ten repeated fivefold held-out projections. (D) Outer-test ROC-AUCs for individual features in the diagnostic panels. (E) Repeat-aggregated ROC curves for the three-feature panel (PON1, HP and C9) and the four-feature panel additionally containing IGHM N209. (F) Outer-test ROC-AUCs for individual features in the two subtype panels. (G) Repeat-aggregated ROC curves for the two subtype panels.

Supervised reproduction of the fixed consensus labels yielded outer-test ROC-AUC values of 0.989, 0.982 and 0.975 for PEP, NGP and early fusion, respectively (**Supplementary Fig. 10A**). To test whether OV subtype structure was reproducible, we performed strict de novo k=2 rediscovery in training folds and projected held-out OV samples (**Fig. 6C**). PEP rediscovery gave projection agreement of 0.932, adjusted Rand index (ARI) 0.744 and normalized mutual information (NMI) 0.667; NGP was more stable (agreement 0.990, ARI 0.960 and NMI 0.939). Held-out cross-omic PEP-NGP agreement reached 0.926 (ARI 0.723 and NMI 0.637). Sample-level assignment frequencies and the stability of feature selection and held-out assignments are shown in **Supplementary Fig. 10B-D**.

Within the nested evaluation, we locked candidate features and separately constructed four concise exploratory panels, two for diagnosis and two for subtype discrimination. (**Fig. 6D-G**). Diagnostic panel A comprising PON1, HP and C9 achieved ROC-AUC 0.833±0.013, which increased to 0.843±0.026 after adding IGHM_N209_N4H6F2 (panel B). Two three-feature subtype panels achieved ROC-AUC 0.997±0.005 (panel C: C2_N621_N2H5, SERPINA3 and IGHG1_N180_N4H5F1) and 0.996±0.006 (panel D: C9, C2_N621_N2H5 and HP_N241_N4H6F1S1), respectively. An independent discovery–validation study of HGSOC reported reduced PON1 and elevated HP, C9, and SERPINA3/AACT signals in lectin-enriched glycoproteins, with C9 alterations also detected in prediagnostic samples (Dutt et al., 2023). Related ovarian cancer studies independently identified altered sialylation of PON1 and HP, increased sialyl-Lewis X on HP and SERPINA3, and elevated C9 and SERPINA3 in serum-derived extracellular vesicles (Lai et al., 2022; Saldova et al., 2007; Shetty et al., 2012). Although PON1, HP, C9, and SERPINA3 were measured in this study as non-glycosylated PEP features, these findings support the biological plausibility of these protein-level observations. Among the subtype-discriminating N-glycopeptides, IGHG1_N180_N4H5F1 was decreased in OV relative to the combined HC and Ben group but remained higher in C1 than in C2. It was previously reported to be reduced in EOC relative to healthy controls in both the discovery and independent validation cohorts(Lai et al., 2022), supporting its disease association, whereas its C1-C2 discrimination extends this observation to within-OV molecular heterogeneity. Representative annotated MS2 spectra and graphical fragmentation maps of C2_N621_N2H5, IGHG1_N180_N4H5F1 and HP_N241_N4H6F1S1 were shown in **Supplementary Figs. 11-13**. Each candidate N-glycopeptide had more than six site-diagnostic ions. HP_N241_N4H6F1S1 was identified in both HGSOC tissue cohorts, In the Hu dataset, it was assigned to the complement/coagulation-enriched IG3 group, although its OV increase was not statistically significant.

## 4. Conclusion

In conclusion, we established a cohort-scale workflow combining matched serum proteomics and N-glycopeproteomics profiling with high-pH reversed-phase C18 fractionation and TMT-labelling, enabling deep characterization of 12,634 intact N-glycopeptides across 100 participants. By integrating differential, cross-sectional disease-state trend and within-cancer subtype evidence, the DTS framework generated an information-rich candidate pool spanning protein abundance and site-specific glycosylation. Although the present data do not establish a highly specific early-detection marker, they identified two internally stable ovarian cancer molecular states that were concordantly detected across peptide and glycopeptide layers. Their distinct sialylation and fucosylation patterns, together with residual glycopeptide signals after carrier-protein adjustment, indicate that ovarian cancer heterogeneity comprises coordinated protein remodelling and a smaller glycosylation-specific component. Leakage-controlled modelling further showed that compact exploratory panels could distinguish these molecular states with high accuracy. Collectively, this study provides an integrated experimental and analytical strategy for elucidating serum N-glycoproteome heterogeneity and prioritizing candidates for ovarian cancer diagnosis and molecular stratification.

## Data availability

The raw LC-MS/MS data will be available through the iProX Consortium (https://www.iprox.cn/page/home.html, (Chen et al., 2022; Ma et al., 2019)) with Project ID IPX0020875000. (share data: https://www.iprox.cn/page/SSV024.html;url=1794920429428uxvK, password: YZ61).

## CRediT authorship contribution statement

**Xiaoyu Hu**: all experiments, data analysis, visualization, draft. **Tianqing Yan**: sample collection, clinical information collection. **Renquan Lu**: project administration. **Zhixin Tian:** project administration, funding acquisition.

## Acknowledgements

This research was financially supported by National Nature Science Foundation of China (22074105, 82503569 and 82373383) and Shanghai Science and Technology Commission (14DZ2261100).

