## Supplementary material for "A differential-trend-subtype strategy for integrated serum proteomics and N-glycoproteomics reveals ovarian cancer-associated remodelling": Supplemental Figures.docx

**Contents**

**Supplementary Figure 1.** Quality control supports consistent feature recovery across the NGP and PEP datasets.

**Supplementary Figure 2.** Sequence context of identified N-glycosylation sites recapitulates the canonical N-X-S/T motif.

**Supplementary Figure 3.** NGP profiling extends the accessible serum concentration range and reduces signal dominance by the most abundant proteins.

**Supplementary Figure 4.** Global profiles of three groups.

**Supplementary Figure 5.** Structural composition and direction-specific glycan preferences of the 68 D&T-shared intact N-glycopeptides.

**Supplementary Figure 6.** Cross-study overlap and compartment-specific directionality of site-specific N-glycopeptides.

**Supplementary Figure 7.** Consensus stability and stage-sensitivity analyses of ovarian-cancer molecular states.

**Supplementary Figure 8.** The structural distribution of N-glycopeptides related to the OV subtype.

**Supplementary Figure 9.** Fold-local DTS prioritization supports compact cross-omic diagnostic and subtype panels.

**Supplementary Figure 10.** Fixed-label reproduction and nested stability of ovarian-cancer subtype assignments.

**Supplementary Figure 11.** Representative annotated MS2 spectrum and graphical fragmentation map (inset) of VVLHPNYSQVDIGLIK carrying N4H6F1S1 at N241 from haptoglobin (HP; P00738).

**Supplementary Figure 12.** Representative annotated MS2 spectrum and graphical fragmentation map (inset) of QSVPAHFVALNGSK carrying N2H5F0S0 at N621 from complement C2 (C2; P06681).

**Supplementary Figure 13.** Representative annotated MS2 spectrum and graphical fragmentation map (inset) of EEQYNSTYR carrying N4H5F1S0 at N180 from immunoglobulin heavy constant gamma 1 (IGHG1; P01857).

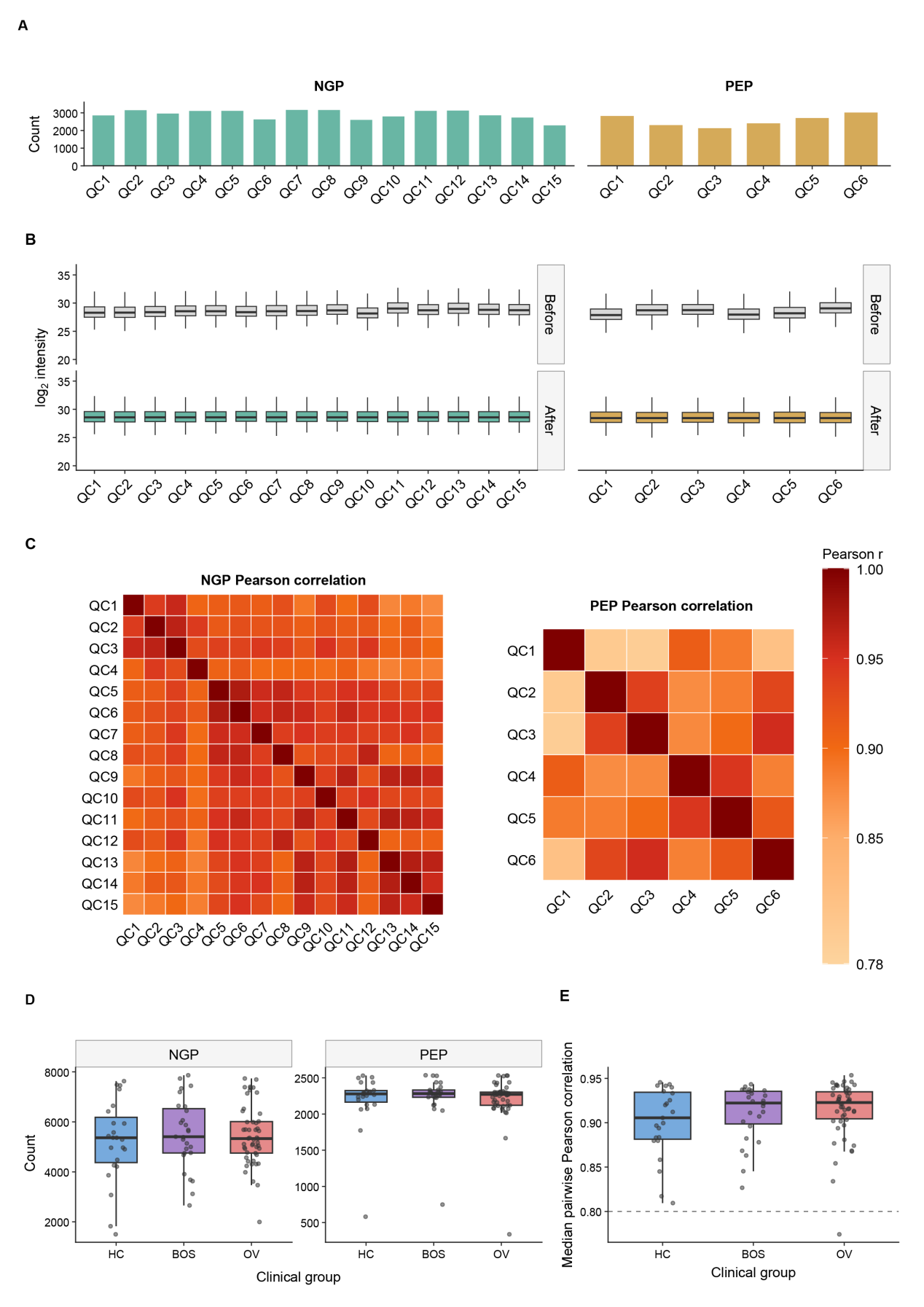
**Supplementary Figure 1. Quality control supports consistent feature recovery across the NGP and PEP datasets.** (A) Numbers of non-missing features in the 15 NGP and 6 PEP technical QC channels. (B) Log2-intensity distributions of filtered QC features before and after median normalization; NGP features quantified in at least 8 of 15 QC channels and PEP features quantified in at least 3 of 6 QC channels were retained. (C) Pairwise Pearson correlation matrices for features complete across all QC channels in each omic layer. (D) Numbers of quantified NGP and PEP features per sample in the HC, Ben and OV groups. (E) Median pairwise Pearson correlations among peptide technical replicates in each clinical group; the dashed line denotes r = 0.80. NGP, intact N-glycopeptide; PEP, peptide; QC, quality control; HC, healthy control; Ben, benign; OV, ovarian cancer.

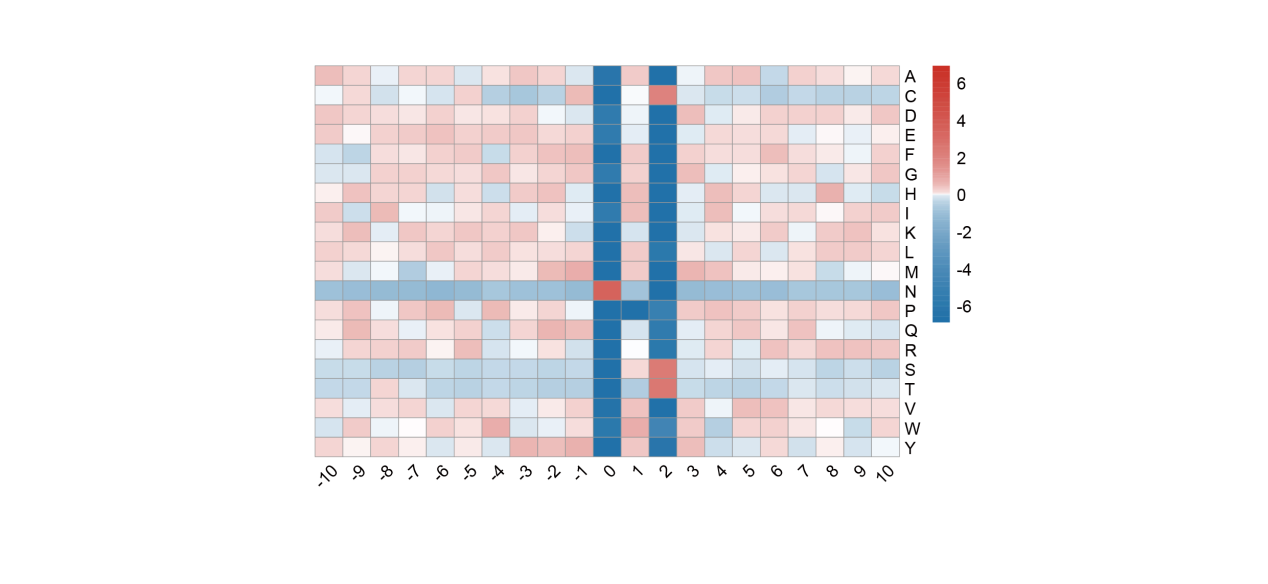

**Supplementary Figure 2. Sequence context of identified N-glycosylation sites recapitulates the canonical N-X-S/T motif.** Heatmap of log2 amino-acid enrichment relative to the global background frequency from positions -10 to +10 around the modified asparagine (position 0).

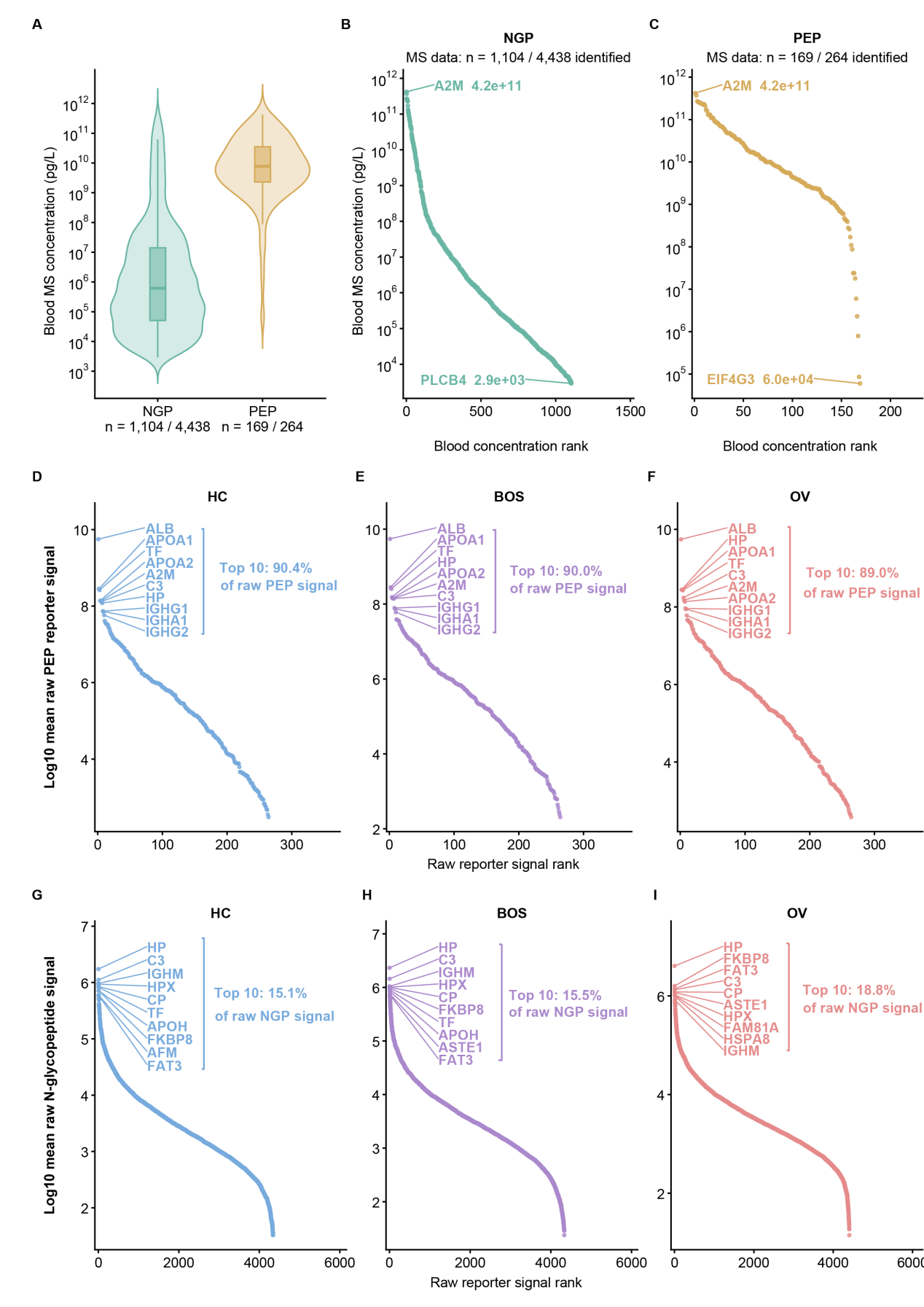

**Supplementary Figure 3. NGP profiling extends the accessible serum concentration range and reduces signal dominance by the most abundant proteins.** (A) Human Protein Atlas blood mass-spectrometry concentrations for proteins identified by NGP or PEP profiling; matched proteins are shown as violin and box plots (NGP, n=1,104; PEP, n=169). (B,C) Ranked blood concentrations for NGP- and PEP-identified proteins, respectively. (D-F) Ranked raw PEP reporter-ion signal in HC, Ben and OV samples; the ten most abundant proteins account for 90.4%, 90.0% and 89.0% of total signal, respectively. (G-I) Ranked raw NGP reporter-ion signal in the same groups; the ten most abundant proteins account for 15.1%, 15.5% and 18.8% of total signal, respectively. Concentrations are in pg·L^-1^.

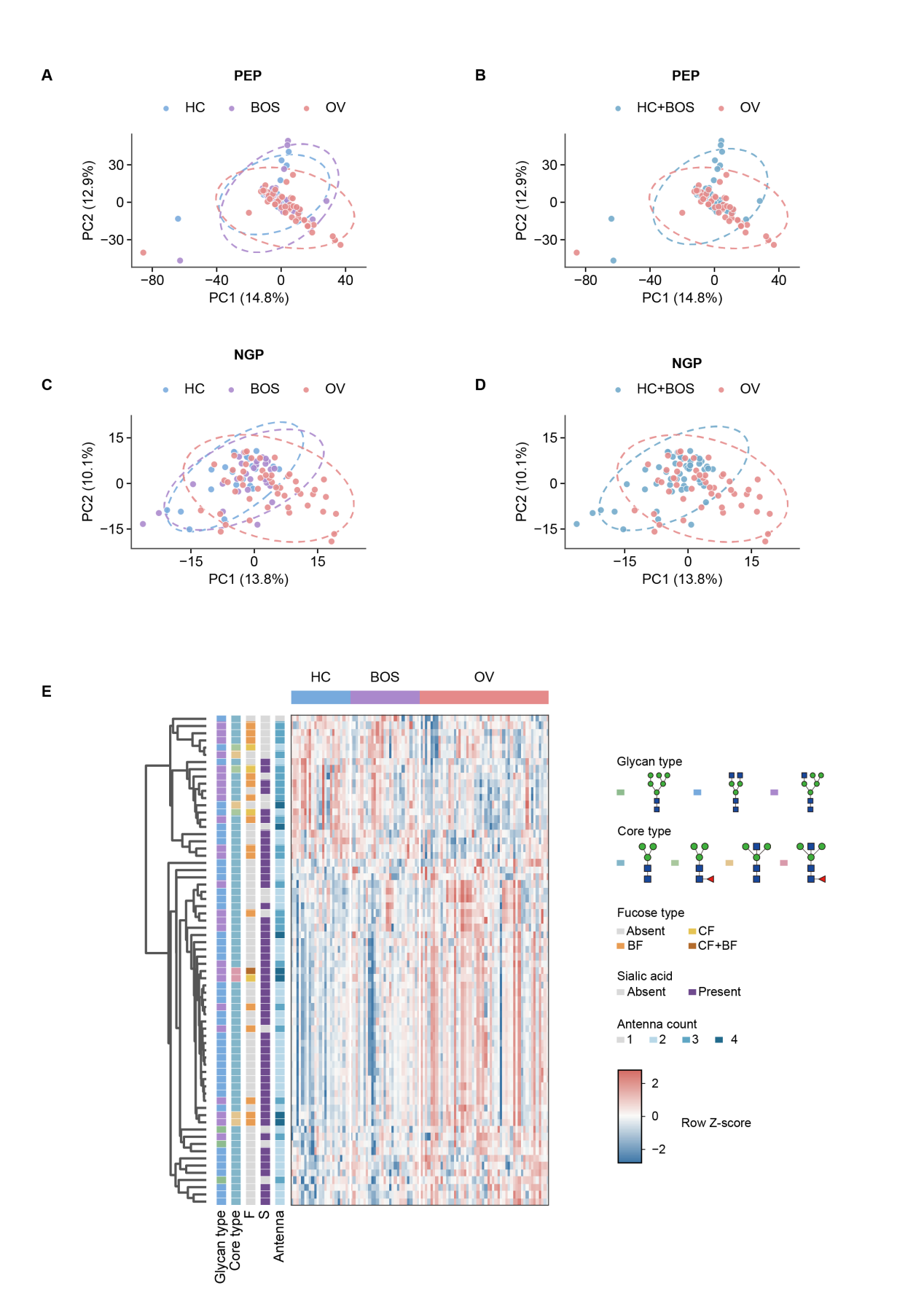

**Supplementary Figure 4. Global profiles of three groups.** (A,B) Principal component analysis (PCA) of 1,763 PEP features quantified in more than 70% of the 100 samples, coloured by HC, Ben and OV group (A) or by HC+Ben and OV status (B). (C,D) PCA of 512 NGP features passing the same completeness criterion, coloured as in A and B. Missing values were completed by k-nearest-neighbour imputation (k=10), and retained features were centred and scaled before PCA. Dashed outlines denote 95% covariance ellipses. (E) Sample-level NGP abundance heatmap across HC, Ben and OV, shown as row Z-scores with annotations for glycan type, core type, fucosylation, sialylation and antenna count.

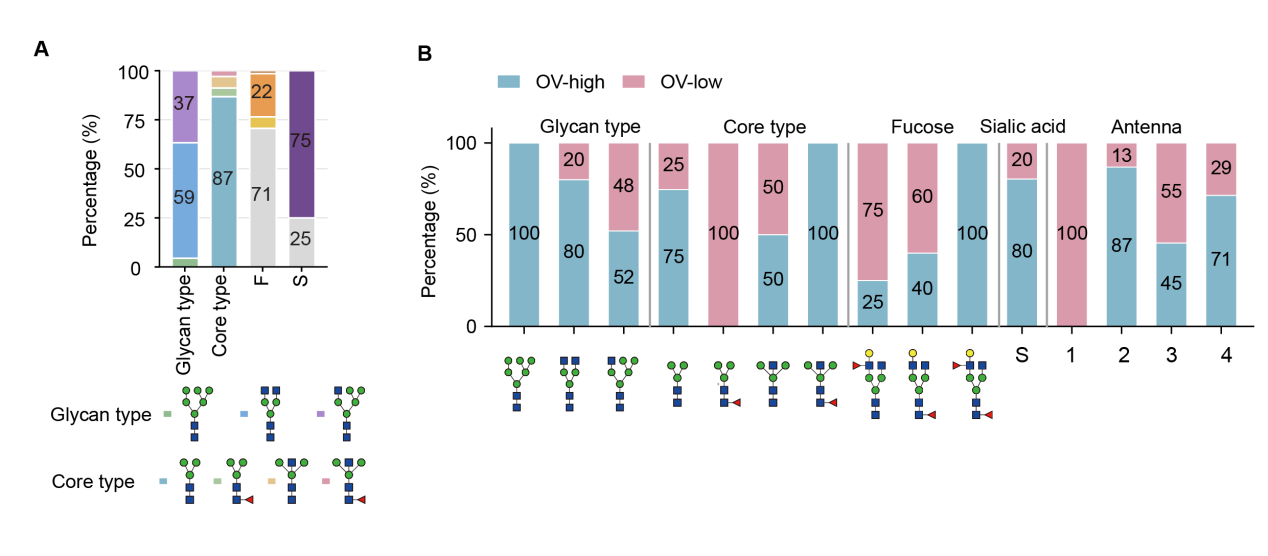

**Supplementary Figure 5. Glycan preferences of the 68 D&T-shared intact N-glycopeptides.** (A) Overall distributions of glycan type, core type, fucosylation and sialylation among the shared NGPs. (B) Percentages of OV-high and OV-low features within each glycan type, core class, fucosylation class, sialylation status and antenna class. OV-high and OV-low groups contained 48 and 20 NGPs, respectively.

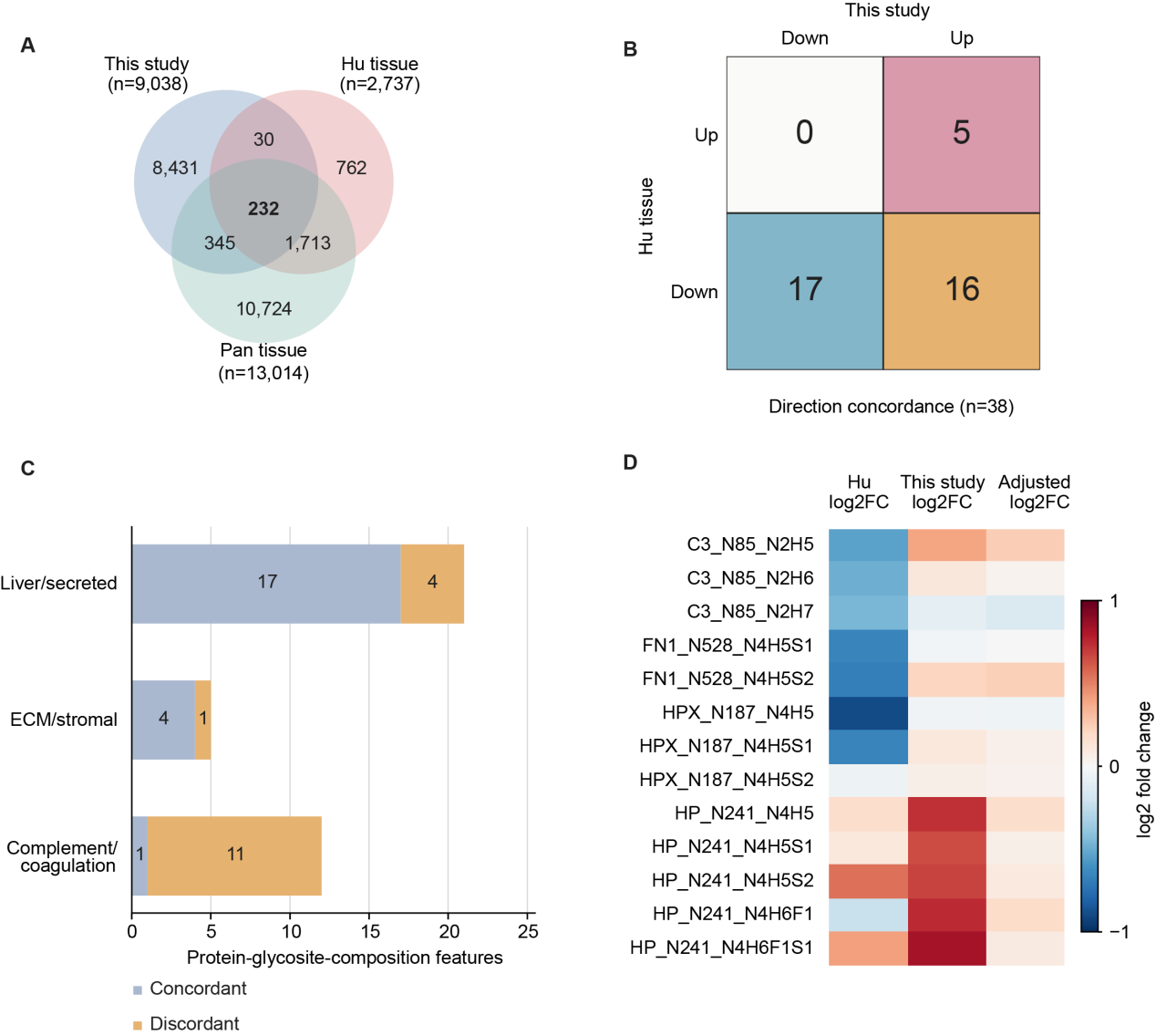

**Supplementary Figure 6. Cross-study overlap and compartment-specific directionality of site-specific N-glycopeptides.** (A) Overlap of protein-glycosite-composition features between this serum study and ovarian-cancer tissue datasets from Hu et al. and Pan et al., matched by gene, sequence-harmonized canonical N-glycosite and N/H/F/S composition. (B) Direction concordance for 38 features with comparable tumour-associated directions in this study and Hu tissue. (C) Concordant and discordant features stratified into liver/secreted, extracellular matrix (ECM)/stromal and complement/coagulation biological-source classes. (D) Hu, serum and carrier-adjusted log2 fold changes for direction-switching glycoforms sharing the same protein and glycosite. Hu values compare tumour with non-tumour tissue; this-study values compare OV with HC+Ben. Carrier-adjusted log2 fold changes account for matched protein abundance.

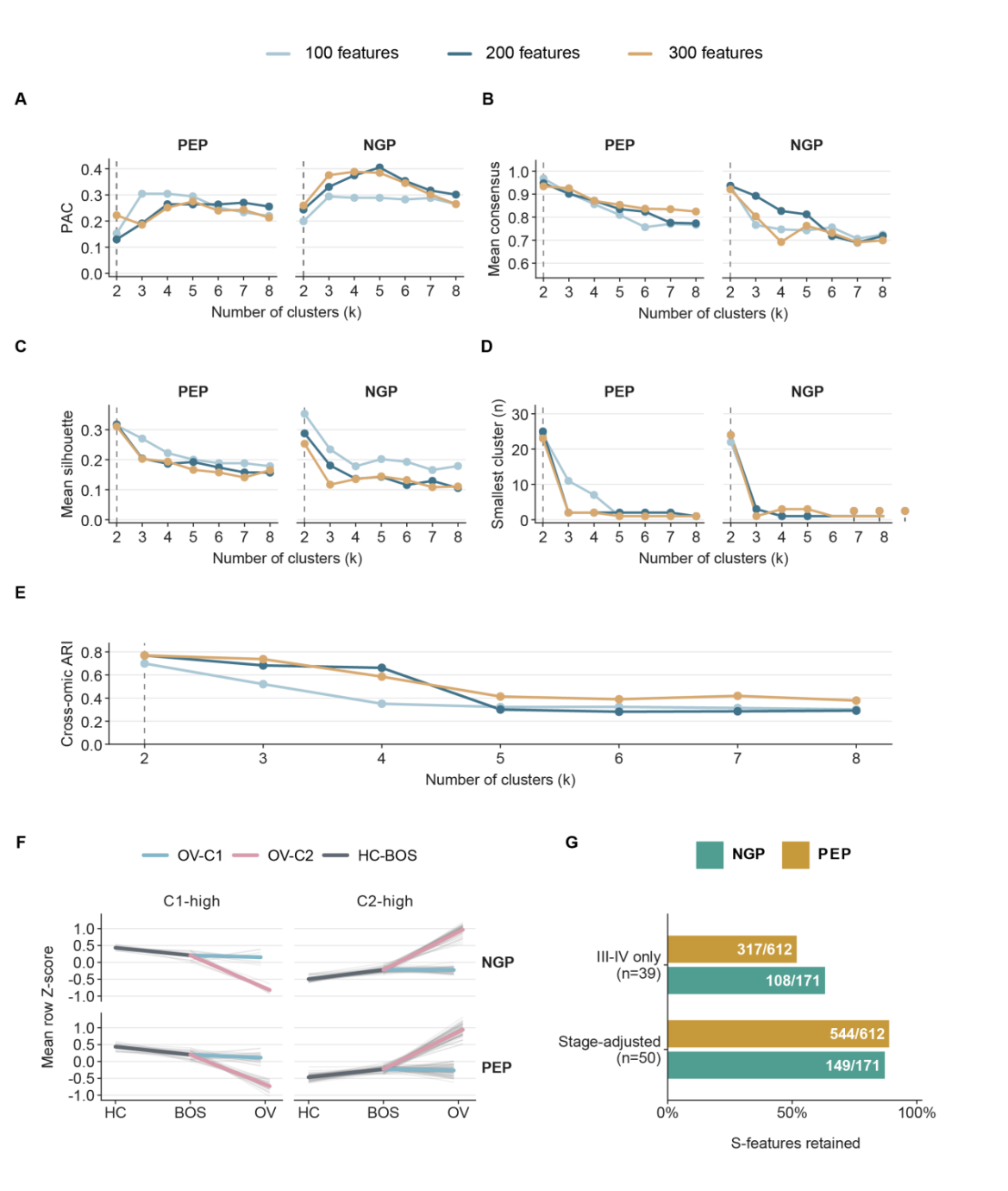

**Supplementary Figure 7. Consensus stability and stage-sensitivity analyses of ovarian-cancer molecular states.** (A-D) Proportion of ambiguous clustering (PAC), mean consensus, mean silhouette width and smallest cluster size for independent PEP and NGP consensus clustering across k=2-8 and the 100, 200 or 300 most variable features. (E) Adjusted Rand index (ARI) between PEP- and NGP-derived clustering results; the dashed line marks k=2 and the prespecified top-200 feature set is highlighted. Consensus clustering used Pearson-correlation distance, average linkage and 500 resamples with 80% sample and feature subsampling. (F) Adjusted group-mean row Z-scores DTS features in PEP (n = 83) and NGP (n = 54), separated into C1-high and C2-high patterns. (G) Numbers and fractions of subtype-associated features retained with the original C1/C2 labels fixed after stage adjustment in all OV samples (PEP, 544 of 612; NGP, 149 of 171; n = 50) or restriction to stage III–IV disease (PEP, 317 of 612; NGP, 108 of 171; n = 39). D, differential; T, trends across HC-Ben-OV; S, subtype.

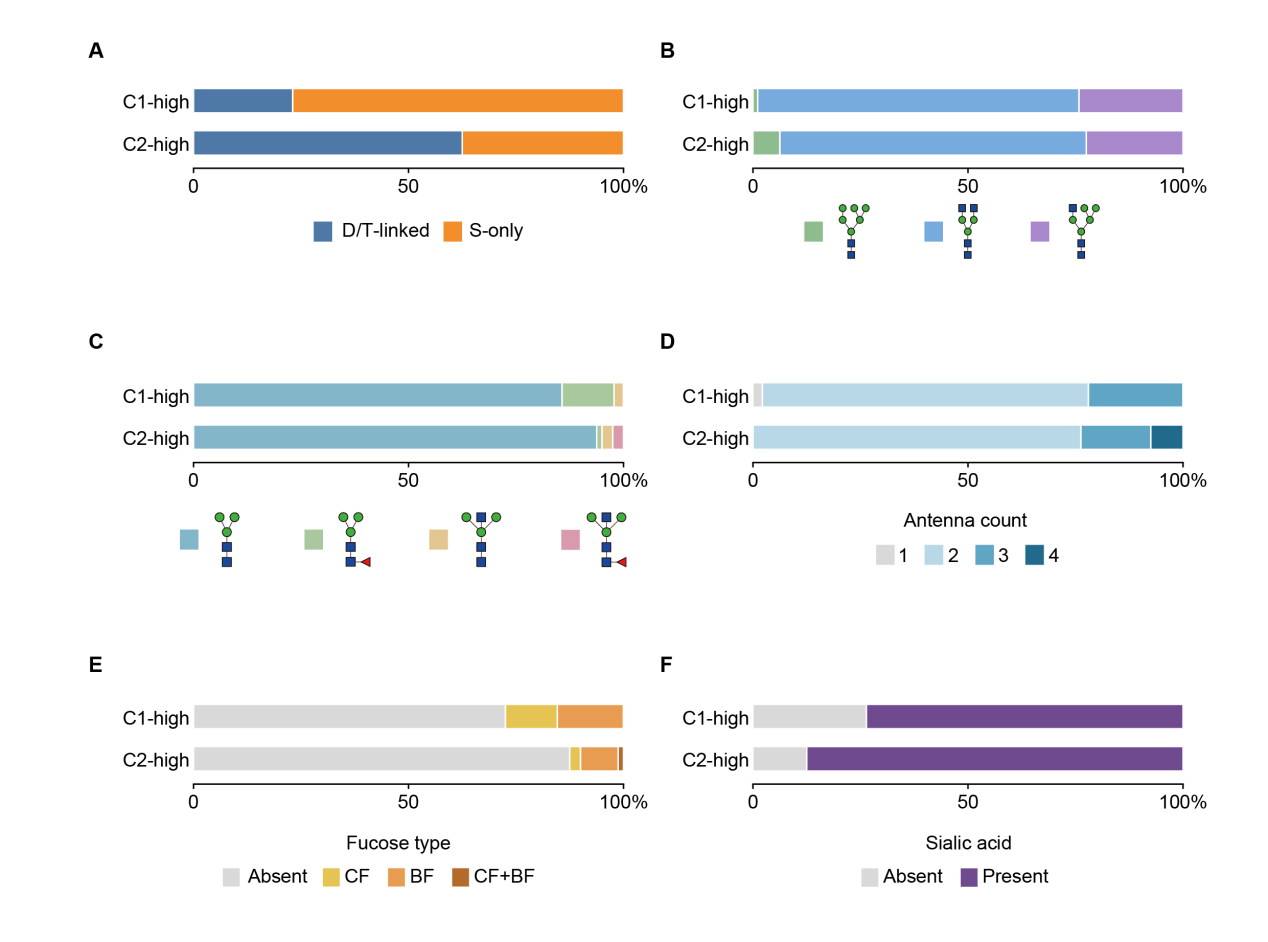

**Supplementary Figure 8. The structural distribution of N-glycopeptides related to the OV subtype.** (A-F) Feature-level distributions of disease-axis relation (A), glycan type (B), core type (C), antenna count (D), fucosylation class (E) and sialylation status (F) among 91 C1-high and 80 C2-high subtype-associated NGPs.

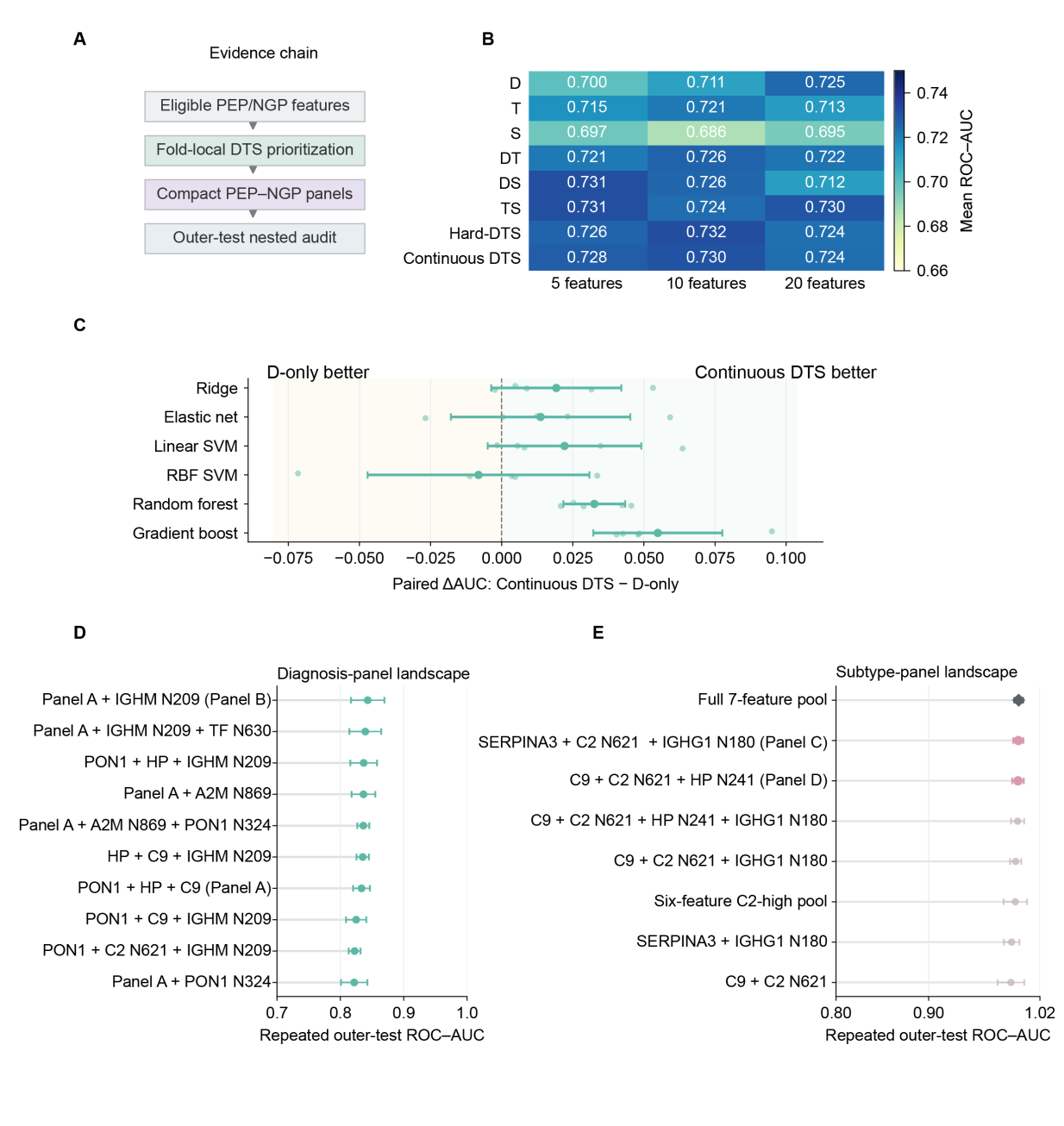

**Supplementary Figure 9. Fold-local DTS prioritization supports compact cross-omic diagnostic and subtype panels.** (A) Analysis workflow from eligible PEP and NGP features through fold-local DTS prioritization, compact panel construction and outer-test evaluation. (B) Mean outer-test ROC-AUC for the D-, T- and S-only strategies, their pairwise combinations, Hard-DTS and Continuous-DTS at panel sizes of 5, 10 and 20 features. (C) Paired outer-test AUC differences between continuous-DTS and D-only ranking across six classifiers. (D) Repeated outer-test ROC-AUCs for candidate HC+Ben versus OV diagnostic panels. (E) Repeated outer-test ROC-AUCs for the full seven-feature subtype candidate pool and compressed panels.

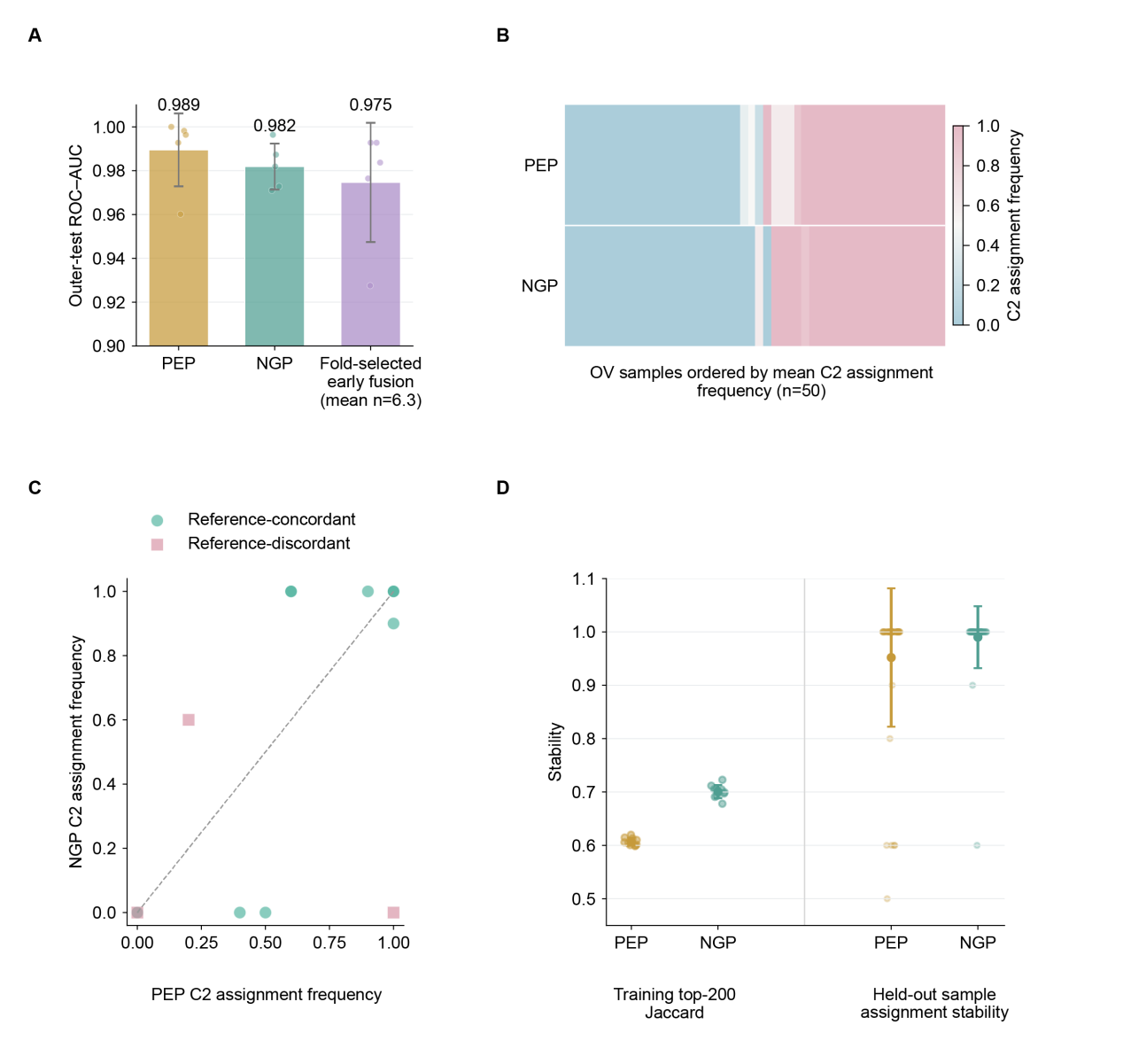

**Supplementary Figure 10. Fixed-label reproduction and nested stability of ovarian-cancer subtype assignments.** (A) Outer-test ROC–AUC for supervised reproduction of the fixed consensus C1/C2 labels using PEP, NGP or fold-selected early-fusion features. (B) C2 assignment frequenciess for each held-out OV sample after training-fold subtype rediscovery and projection in the PEP and NGP feature spaces. (C) Comparison of PEP and NGP C2 assignment frequencies. (D) Stability of the training top-200 feature sets, measured by the Jaccard index, and stability of held-out sample assignments for PEP and NGP.

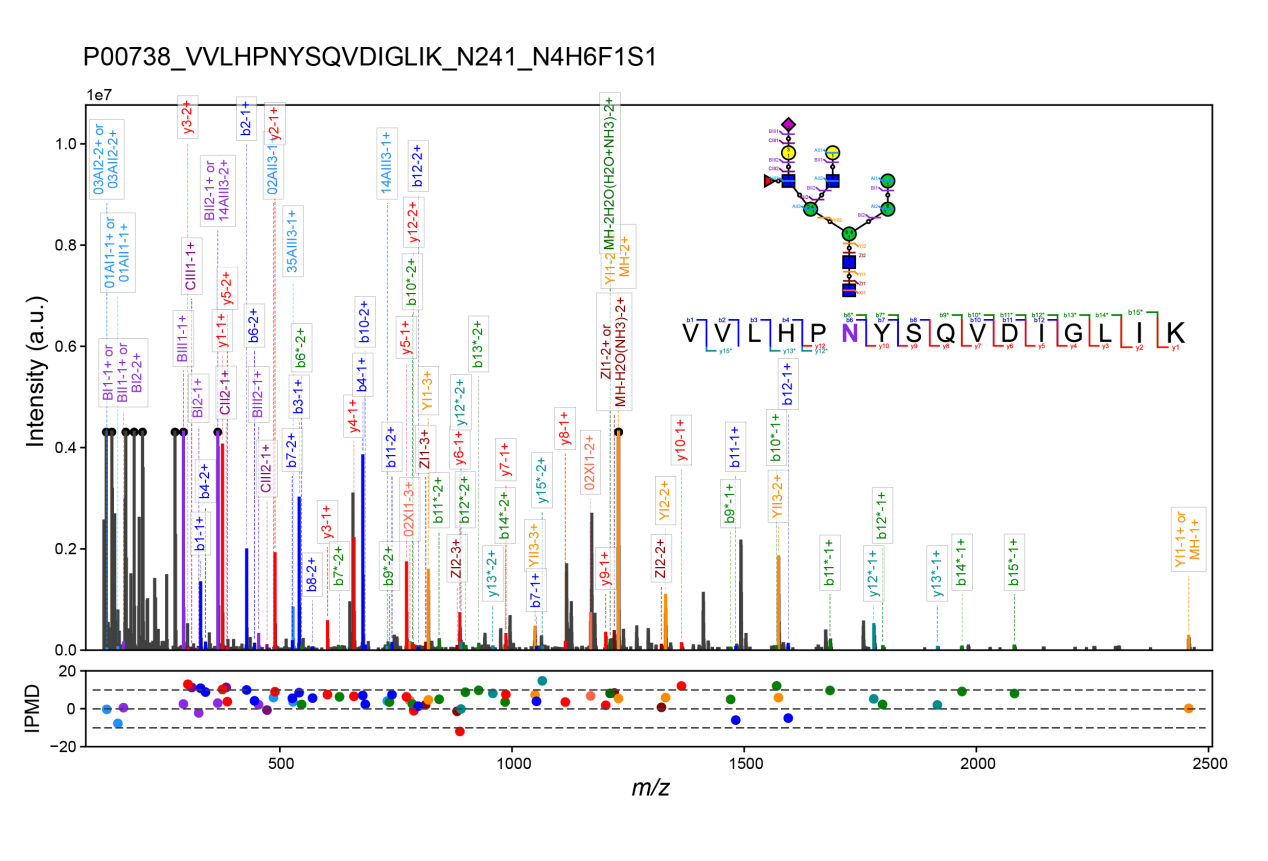

**Supplementary Figure 11. Representative annotated MS2 spectrum and graphical fragmentation map (inset) of VVLHPNYSQVDIGLIK carrying N4H6F1S1 at N241 from haptoglobin (HP; P00738).** The lower panel shows isotopic-pattern mass deviations (IPMDs) for assigned fragment ions.

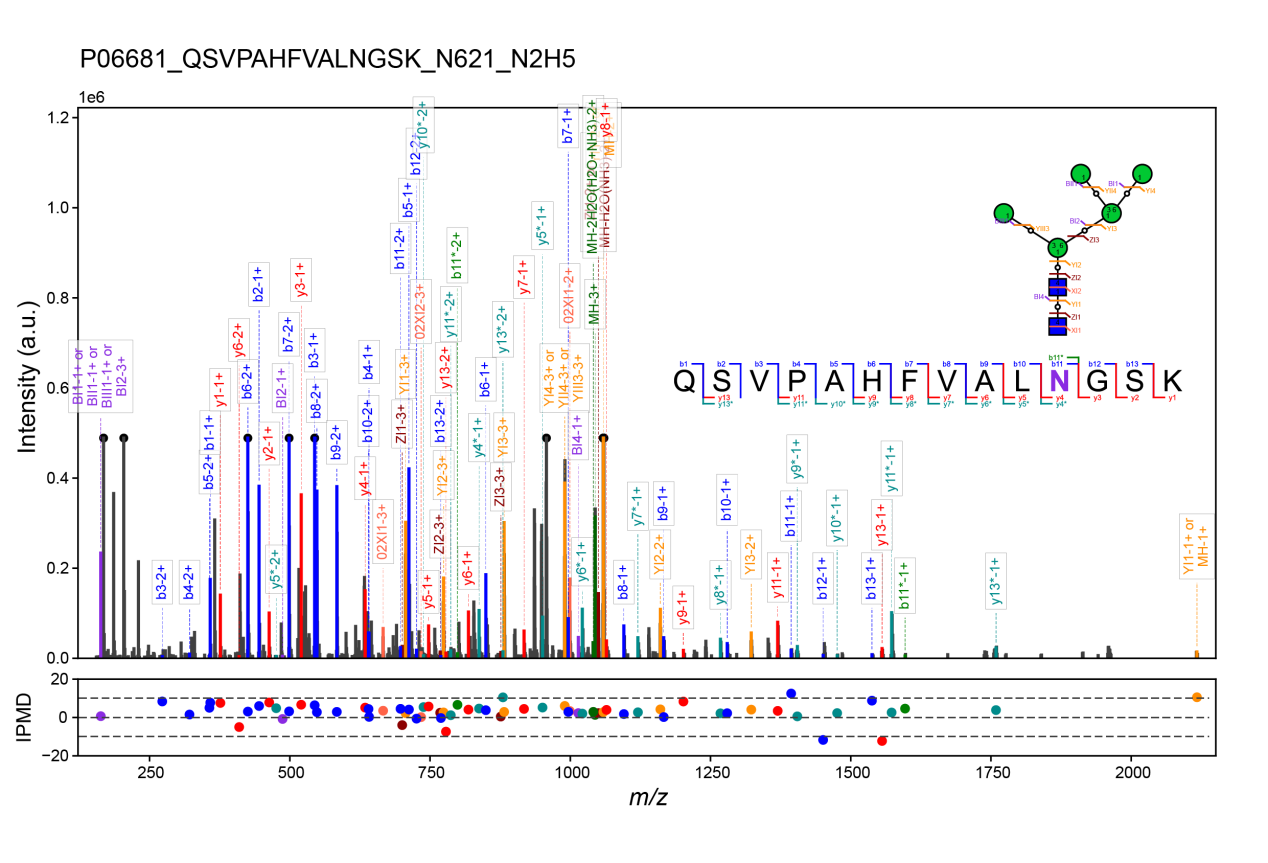
**Supplementary Figure 12. Representative annotated MS2 spectrum and graphical fragmentation map (inset) of QSVPAHFVALNGSK carrying N2H5F0S0 at N621 from complement C2 (C2; P06681).** The lower panel shows IPMDs for assigned fragment ions.

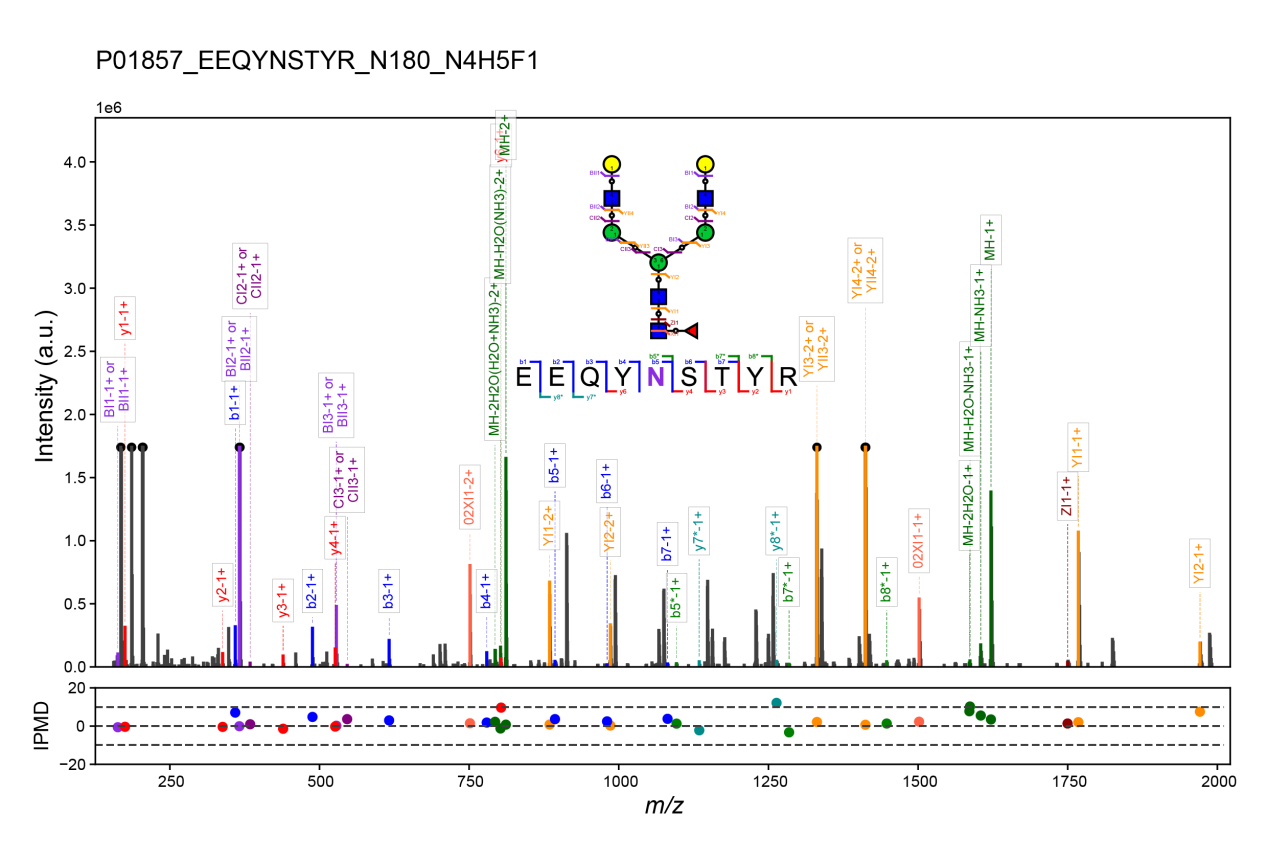

**Supplementary Figure 13. Representative annotated MS2 spectrum and graphical fragmentation map (inset) of EEQYNSTYR carrying N4H5F1S0 at N180 from immunoglobulin heavy constant gamma 1 (IGHG1; P01857).** The lower panel shows IPMDs for assigned fragment ions.
